# The Mitochondrial Calcium Uniporter of Pulmonary Type 2 Cells Determines Severity of ARDS

**DOI:** 10.1101/2021.01.18.427173

**Authors:** Mohammad Naimul Islam, Galina A. Gusarova, Shonit R. Das, Li Li, Eiji Monma, Murari Anjaneyulu, Edward Owusu-Ansah, Sunita Bhattacharya, Jahar Bhattacharya

## Abstract

Acute lung immunity to inhaled pathogens elicits defensive pneumonitis that may convert to the Acute Respiratory Distress Syndrome (ARDS), causing high mortality. Mechanisms underlying the conversion are not understood, but are of intense interest because of the ARDS-induced mortality in the ongoing Covid-19 pandemic. Here, by optical imaging of live lungs we show that key to the lethality is the functional status of mitochondrial Ca^2+^ buffering across the mitochondrial Ca^2+^ uniporter (MCU) in the lung’s alveolar type 2 cells (AT2), which protect alveolar stability. In mice subjected to ARDS by airway exposure to lipopolysaccharide (LPS), or to *Pseudomonas aeruginosa*, there was marked loss of MCU expression in AT2. The ability of mice to survive ARDS depended on the extent to which the MCU expression recovered, indicating that the viability of Ca^2+^ buffering by AT2 mitochondria critically determines ARDS severity. Mitochondrial transfer to enhance AT2 MCU expression might protect against ARDS.

## INTRODUCTION

The acute immune response at environment-facing epithelial barriers provides rapid systemic protection against ambient pathogens. The lung, which is particularly prone to challenge by inhaled pathogens, develops brisk immunity to clear the pathogens and protect barrier properties of the alveolar epithelium, the site of blood oxygenation. Pneumonia due to non-resolving inflammation causes barrier injury and pulmonary edema, impairing oxygenation, and setting the stage for the Acute Respiratory Distress Syndrome (ARDS), a condition associated with high mortality (Bellani et al., 2016). Mechanistic understanding is lacking as to how benign pneumonias lead to life-threatening ARDS.

The mortality risk in pneumonia is due not only to the infectious load of the inhaled organism, but also to the injurious host response that involves failure of the alveolar epithelial fluid barrier, which causes the characteristic pulmonary edema of ARDS (Bhattacharya and Matthay, 2013). The epithelium consists of structural type 1 cells that form the alveolar wall, and the metabolically significant type 2 cells (hereafter, AT2). AT2 are particularly important for barrier maintenance, since they secrete surfactant that stabilizes alveolar patency and protect against pulmonary edema (Mora et al., 2000; Wu et al., 2020). However, immunity-induced mechanisms that impair the secretion are not well understood.

AT2 mitochondria are important in this regard, since they provide ATP to support surfactant secretion (Islam et al., 2014). Although not shown directly for AT2, mitochondrial calcium (mCa^2+^) buffering modulates the cytosolic Ca^2+^ (cCa^2+^), unchecked increases in which cause barrier weakening (Hough et al., 2019). Indirect support for a mitochondrial role in lung immune injury comes from findings that replenishment of the immune-challenged alveolar epithelium with exogenous mitochondria protects against lung injury (Islam et al., 2012). However, the role of mitochondrial buffering in this protection remains unclear.

Mitochondrial buffering involves Ca^2+^ flux from the cytosol to the mitochondrial matrix across the mitochondrial calcium uniporter (MCU) (Baughman et al., 2011; De Stefani et al., 2011). The flux is regulated by MCU-associated proteins (Kamer and Mootha, 2015), as well as by oxidative (Dong et al., 2017) and transcriptional (Favaro et al., 2019; Shanmughapriya et al., 2015) mechanisms. Studies in mice involving global (Pan et al., 2013) or tissue-specific *MCU* deletion (Kwong et al., 2015), or overexpression of dominant-negative *MCU* (Rasmussen et al., 2015) reveal a determining role of the MCU in multiple aspects of organ function, inlcuding body size determination (Pan et al., 2013), exercise tolerance (Gherardi et al., 2019; Kwong et al., 2018; Pan et al., 2013), myocardieal infarction (Kwong et al., 2015; Luongo et al., 2015), pulmonary fibrosis (Gu et al., 2019), insulin secretion (Georgiadou et al., 2020), fibroblast differentiation (Lombardi et al., 2019), and hepatic lipidosis (Tomar et al., 2019). Despite this extensive evidence for MCU involvement in organ function, the reported data rely on *a priori* genetic modifications that do not replicate natural pathogenesis. Understanding is therefore lacking as to whether disease processes modify the MCU to exacerbate pathophysiology.

Here, we address this issue in LPS and bacterial models of lung infection, using a novel *in situ* assay of mitochondrial buffering in AT2 of live alveoli by means of real-time confocal microscopy (Islam et al., 2012). We could thereby, determine the first dynamic Ca^2+^ responses in the cytosol and mitochondria in relation to the progression of acute immunity. Our findings indicate that by the first post-infection day, there was marked depletion of both message and protein levels of AT2 MCU, together with loss of buffering. Unexpectedly, sustained MCU depletion promoted mortality, which could be mitigated by MCU repletion, revealing for the first time the critical role of AT2 MCU in immune alveolar injury.

## RESULTS

### The MCU and alveolar homeostasis

For *in situ* assay of MCU function in AT2, we quantified mitochondrial buffering and its effect on surfactant secretion. We identified AT2 as cells that stained for surfactant-containing lamellar bodies (LBs) and for surfactant protein B (Figure S1A) ((Islam et al., 2014). By live confocal microscopy in conjunction with alveolar microinfusion, we loaded the alveolar epithelium with the dyes, rhod2 and fluo4 to quantify mCa^2+^ and cCa^2+^, respectively. To confirm compartmental distribution of the dyes, we gave alveolar microinfusion of a mild detergent, which released fluo4, but not rhod2 from the epithelium, affirming that fluo4 was localized to the cytosol and rhod2 to the mitochondrial matrix (Ichimura et al., 2003) (Figure S1B,C). Alveolar stretch due to a 15-second lung hyperinflation induced transient cCa^2+^ oscillations in AT2, but no increase of mean cCa^2+^ (Figure 1A,B). Concomitantly, both the mean and the oscillation amplitude of mitochondrial Ca^2+^ (mCa^2+^) increased (Figure 1A,B and Supplemental video 1). These responses affirmed that the mitochondrial buffering in AT2 protected against increases of the mean cCa^2+^.

**Figure 1.**
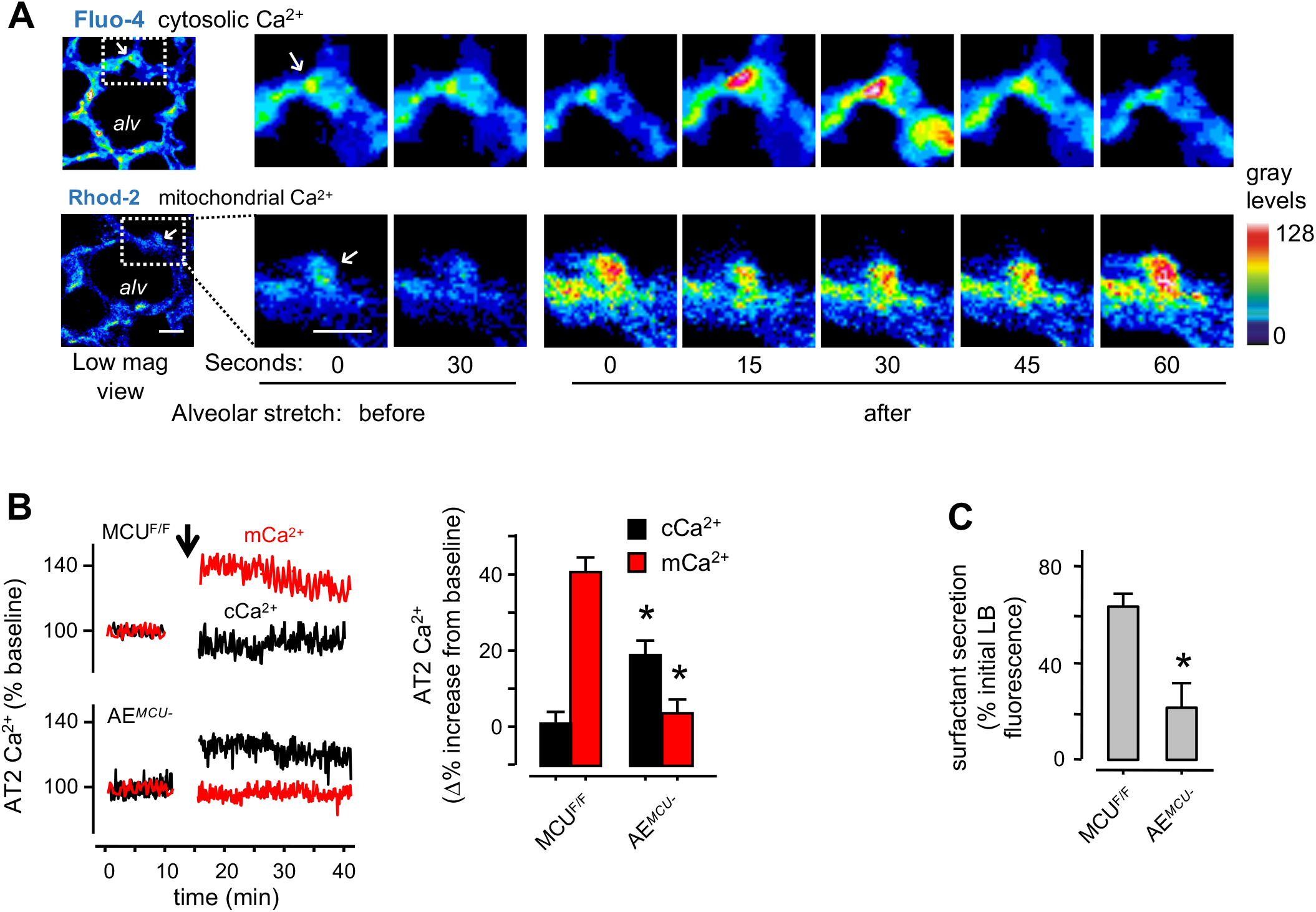
*In situ* AT2 mitochondrial responses to alveolar stretch. (A) Confocal images of a live alveolus (*alv*) show AT2 (*arrows*) in a septum of the alveolar wall, as identified by Lysotracker Red (LTR) staining (not shown). Images show time dependent Ca^2+^ responses in the cytosol (cCa^2+^) and mitochondria (mCa^2+^) of selected AT2 to a single 15-second alveolar stretch induced by increasing airway pressure from 5 to 15 cmH_2_O. Ca^2+^ dyes were given by alveolar microinfusion. Ca^2+^ increases are denoted by increases of pseudocolored gray levels. Images were obtained at airway pressure of 5 cmH_2_O. Scale bar, 10μm. (B) Tracings from an experiment and group data show AT2 Ca^2+^ responses to alveolar stretch (*arrow*). MCU^F/F^, mice floxed for the MCU; AE^*MCU-*^, mice lacking the MCU in the alveolar epithelium. n=4 mice per group, *p<0.05 versus MCU^F/F^. (C) Group data show stretch-induced surfactant secretionas quantified in terms of loss of AT2 fluorescence of lysotracker red (LTR). n=4 lungs for each bar, *p<0.05 versus MCU^F/F^. Group data are mean±SEM.

We tested these responses in MCU^F/F^:SPC-cre mice (AE^*MCU*-^) in which we induced by pre-partum cre recombination in the alveolar epithelium (AE) (Perl et al., 2002; Westphalen et al., 2014) (Figure S1D). We evaluated MCU function in AT2, since mitochondrial function in these cells determines surfactant secretion (Islam et al., 2012). The stretch-induced mitochondrial responses were inhibited pharmacologically by alveolar treatment with the MCU blocker, ruthenium red and they were absent in AE^*MCU-*^ mice (Figures S1E and 1B), confirming the MCU role in the buffering. Further, loss of buffering caused the expected increase of mean cCa^2+^ (Figure 1B). The inhibitor of store Ca^2+^ release, xestospongin C blocked the all Ca^2+^ increases (Figure S1E), consistent with the notion that MCU-dependent mCa^2+^ increases were due to mitochondrial entry of store-released Ca^2+^. Surfactant secretion in intact alveoli is assayed by labeling LBs in AT2, then detecting the time-dependent loss of the fluorescence to a secretion stimulus such as alveolar stretch (Islam et al., 2012; Islam et al., 2014) (Figures S1F-H and 1C). However, this response was absent in AE^*MCU-*^ mice (Figure 1C), as also in mice treated with ruthenium red, indicating that blocking mitochondrial Ca^2+^ entry blocked surfactant secretion (Figure S1G,H). These findings indicated that the AT2 MCU is a major determinant of surfactant secretion, hence of alveolar homeostasis.

### Lung immunity causes MCU loss

Since it is not known whether the acute immune response modifies MCU expression, we determined time dependent responses of AT2 mitochondria in the LPS model of pneumonitis at different doses of intranasal LPS instillation (Islam et al., 2012; Westphalen et al., 2014). LPS doses for different mouse strains are described in Table 2. An intranasal sub-lethal dose of LPS caused marked increase of leukocyte counts in the bronchioalveolar lavage (BAL) within Day 1 (Figure S2A). This brisk immune response recovered to baseline by Day 3 (Figure S2A). Alveolar homeostasis was destabilized on Day1 as indicated by abrogation of surfactant secretion, and increase of extravascular lung water, a measure of pulmonary edema (Figure S2B). Thus, the LPS-induced alveolar injury was transient, providing an opportunity to evaluate early and late mitochondrial responses.

In lungs derived from these LPS-treated mice, the AT2 physiological responses, namely mCa^2+^ increase and surfactant secretion, were present at 4h post-LPS, but not at 24h (Figure 2A,B), indicating that a single instillation of LPS deteriorated alveolar mitochondrial buffering by Day 1. To rule out the possibility that alveolar inflammation may have interfered with alveolar expansion, thereby non-specifically blocking stretch-induced effects, we applied an optogenetic approach to increase cCa^2+^ (Lin et al., 2009). We expressed the light sensitive, cation channel, channelrhodhopsin-2 (ChR2) in the alveolar epithelium (Figure S2C). Excitation of ChR2 by blue light increased AT2 mitochondrial Ca^2+^ in PBS-, but not in LPS-treated lungs (Figures S2D,E and 2C). Hence, the stretch and optogenetics approaches together denoted loss of MCU-induced buffering in AT2 following LPS treatment.

**Figure 2.**
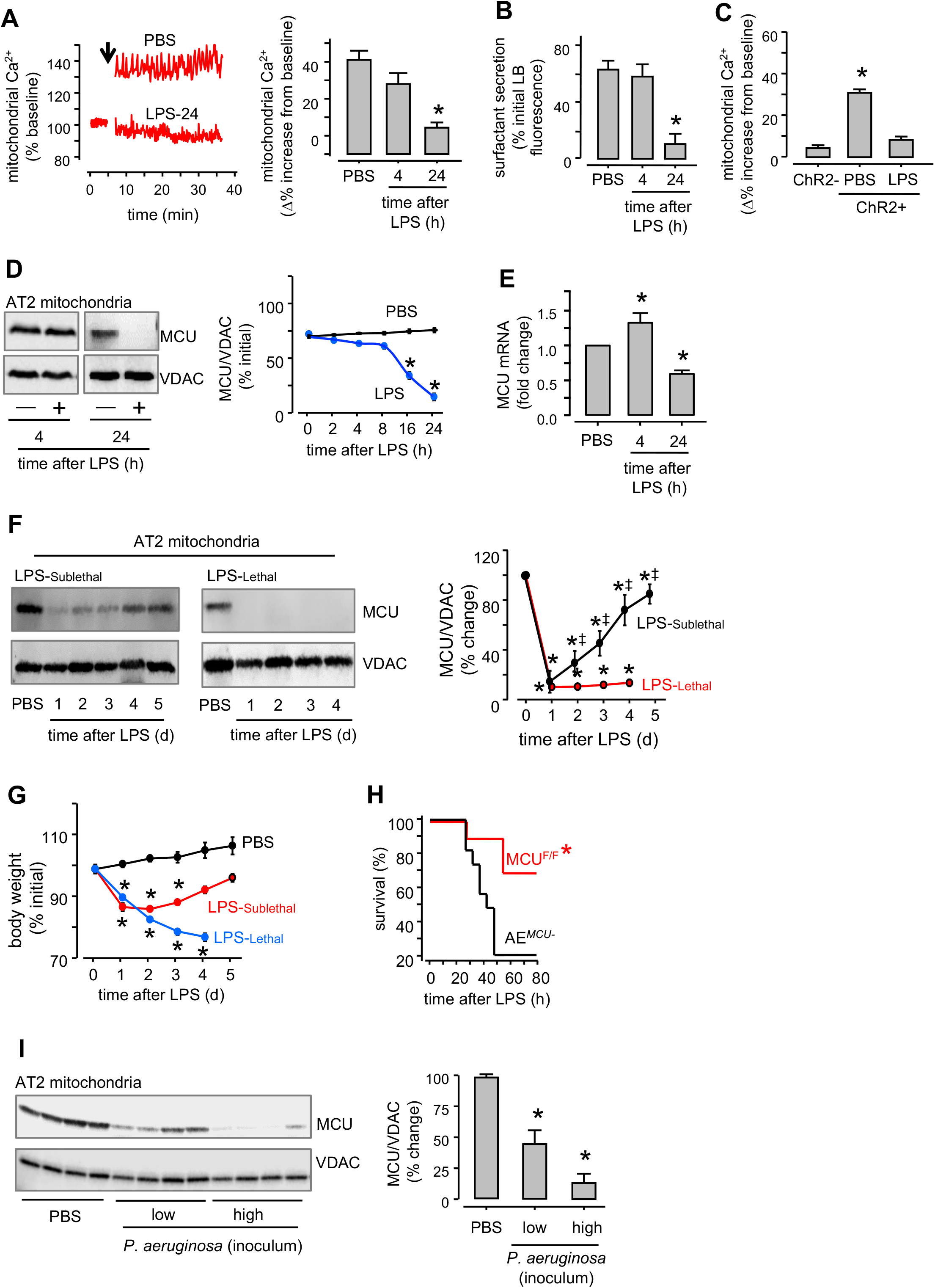
LPS effects on AT2 mitochondria. (A) Tracings from a single experiment and group data show stretch (*arrow*)-induced AT2 mitochondrial responses following the indicated intranasal instillations. LPS was given at sublethal dose (Table 2). *LPS-24*, 24h after intranasal LPS. n=4 lungs per group, *p<0.05 versus PBS. (B) Group data show quantifications of stretch-induced surfactant secretion for the indicated groups. n=4 lungs each bar, *p<0.05 versus PBS. (C) Group data show Ca^2+^ responses of AT2 mitochondria in live alveoli to a 10-second channelrhodopsin (*ChR2*) activation. *ChR2-GFP* plasmid was given by intranasal instillation in liposomal complexes. Lungs were excised 48h after instillation. *ChR2*+ and *ChR2*-are respectively, data for AT2 that were positive or negative for GFP fluorescence in the same lung. n= 4 lungs per group. *p<0.05 versus ChR2-. (D) Immunoblots are shown for mitochondrial calcium uniporter (MCU) in AT2 in mitochondria derived 24h after indicated treatments. LPS was instilled at a sublethal dose in Swiss Webster mice. *VDAC*, voltage dependent anion channel. Tracings show densitometry quantification of MCU/VDAC ratio. Replicated 4 times. *p<0.05 versus PBS. (E) Data are quantifications of the MCU RNA in freshly isolated AT2 from mice given the indicated intranasal instillations. Lungs were excised and mRNA extracted at indicated time points after instillations of sublethal LPS. RNA values were normalized against actin. n=4 lungs for each bar, *p<0.05 versus PBS. (F) MCU immunoblots of AT2 mitochondria and densitometric quantification of MCU/VDAC ratio after intranasal instillations of sublethal (LPS-Sublethal, *left)* and lethal (LPS-Lethal, *right)* LPS in Swiss Webster mice. PBS was instilled on Day 0. Table 2 gives details of LPS doses. PBS. n=3 mice for each bar, *p<0.05 versus PBS; ‡p<0.05 versus post-LPS day 1. (G) Group data show changes in body weight at indicated time points following intranasal PBS (*black*), sublethal (LPS-Sublethal, *red*) and lethal (LPS-Lethal, *blue*) LPS instillations in Swiss Webster mice. n=4 mice for each bar, *p<0.05 versus PBS. (H) Kaplan-Meier plots for mouse survival are for LPS instillations at a lethal dose in mice lacking *MCU* in the alveolar epithelium (*AE^MCU-^*) and in *MCU*-floxed littermates (*MCU^F/F^*). n=10 mice in each group, *p<0.05 versus AE^MCU-^. (I) Immunoblots are shown for the MCU in AT2 mitochondria derived 24h after indicated instillations. *P. aeruginosa* was intranasally instilled in two doses, 1×10^5^ (low inoculum) and 1×10^6^ (high inoculum) colony-forming units (CFU). Group data show densitometry quantification of MCU/VDAC ratio. *p<0.05 versus PBS. Group data are mean±SEM.

To determine whether the loss of buffering reflected changes in MCU expression, we carried out immunoblots on mitochondria isolated from extracted AT2. MCU protein levels were unchanged from baseline for upto 8h post-LPS. Subsequently, the expression progressively decreased to 25% of baseline by 16h and was undetectable by 24h (Figure 2D). Further, as compared with corresponding PBS lungs, MCU RNA was higher in lungs extracted 4h after LPS, than in those extracted at 24h (Figure 2E). However, despite the MCU loss at Day 1 post-LPS, the mitochondrial outer membrane proteins, TOM-20 and VDAC, and the matrix protein HSP60 were well expressed (Figure S2F,G). The proteins of the mitochondrial electron transport chain (ETC) were also well expressed and their activities were not diminished (except for a small decrease in Complex II activity) (Figure S2H,I). Therefore, the MCU loss was not due to a non-specific loss of mitochondrial proteins.

The mitochondrial membrane potential informs mitochondrial fitness. Mitochondrial depolarization stabilizes PINK1 on the outer membrane. PINK1 recruits the E3-ubiquitin ligase, parkin, which initiates mitophagy (Matsuda et al., 2010). To determine whether the MCU impacted mitochondrial potential, we loaded the alveolar epithelium with the potentiometric dye, TMRE, fluorescence of which decreases with mitochondrial depolarization (Hough et al., 2019; Ichimura et al., 2003). Our findings indicate that despite the loss of the MCU, TMRE fluorescence was unchanged (Figure S2J,K), indicating that the AT2 mitochondria were not depolarized, hence they were unlikely to be mitophagy targets of the PINK1-parkin mechanism. To test this hypothesis, we carried out immunoblots on mitochondria derived from AT2 24h after LPS treatment. These studies failed to reveal association of parkin (Figure S2L), suggesting that the MCU loss did not induce a mitophagy signal.

To evaluate the effect of inflammation severity, we exposed mice to intranasal LPS at sublethal or lethal doses (Table 2) (Islam et al., 2012). For the sublethal dose, the MCU loss on Day1 was followed by recovery of MCU expression (Figure 2F). By contrast, for the lethal dose the expression did not recover (Figure 2F). Similarly, although both doses caused loss of body weight by Day 1, the weight recovered for the sublethal, but not the lethal dose (Figure 2G). These findings suggested that adequacy of AT2 MCU expression was a determinant of survival. To further evaluate this question, we instilled LPS in AE^*MCU-*^ mice. Unexpectedly, mortality was markedly higher in AE^*MCU-*^ mice than in littermate floxed controls (Figure 2H), indicating that loss of MCU function in AT2 mitochondria was of systemic significance. Taking our findings together, we interpret that loss of buffering due to MCU depletion in AT2 resulted in failed surfactant secretion, the systemic consequence of which was increased mortality.

Since the effects of purified LPS might not be entirely representative of those induced by Gram-negative bacteria, we infected lungs by intranasal instillation of *Pseudomonas aeruginosa* (PA) at a high and a low dose to induce dose dependent increases of leukocyte counts in the bronchioalveolar lavage by Day 1 (Figure S2M). Concomitantly, the loss of AT2 MCU expression was also dose dependent (Figure 2I). These findings indicated that the lung’s immune response caused a dose dependent loss of AT2 MCU expression.

### LPS-induced mitochondrial H_2_O_2_ degrades MCU of AT2 mitochondria

To determine the role of mitochondrial H_2_O_2_ production in these responses (Carneiro et al., 2018; Ip et al., 2017), we expressed the matrix-targeted, H_2_O_2_ sensor roGFP (Waypa et al., 2010). Transfection of roGFP by intranasal delivery of liposomal plasmid avoids potential difficulties due to mitochondrial uptake of diffusible H_2_O_2_ sensing dyes (Polster et al., 2014). Intranasal LPS instillation progressively increased H_2_O_2_ in AT2 mitochondria (Figures S3A and 3A). To block this response, we expressed catalase in AT2 mitochondria by crossing inducible *SPC-Cre* mice with mice expressing mitochondria-targeted human *catalase* downstream of a floxed STOP codon (Schriner et al., 2005). Post-partum induction of *SPC-Cre* caused catalase expression in AT2 mitochondria of these mice (AT2^CAT^), as confirmed by alveolar immunofluorescence and immunoblot (Figure S3B,C). LPS treatment of AT2^CAT^ mice failed to increase mitochondrial H_2_O_2_, or induce loss of AT2 MCU (Figure 3B,C). Accordingly, MCU function was retained (Figure 3D,E), alveolar inflammation was mitigated (Figure 3F), and mortality was reduced (Figure 3G). Mice pre-treated with the mitochondria-specific anti-oxidant, MitoQ were also protected from the MCU loss (Figures S3D), indicating efficacy of pharmacalogic intervention. Together, these findings indicated that LPS-induced increase in AT2 mitochondrial H_2_O_2_ was a critical mediator of the MCU loss.

**Figure 3.**
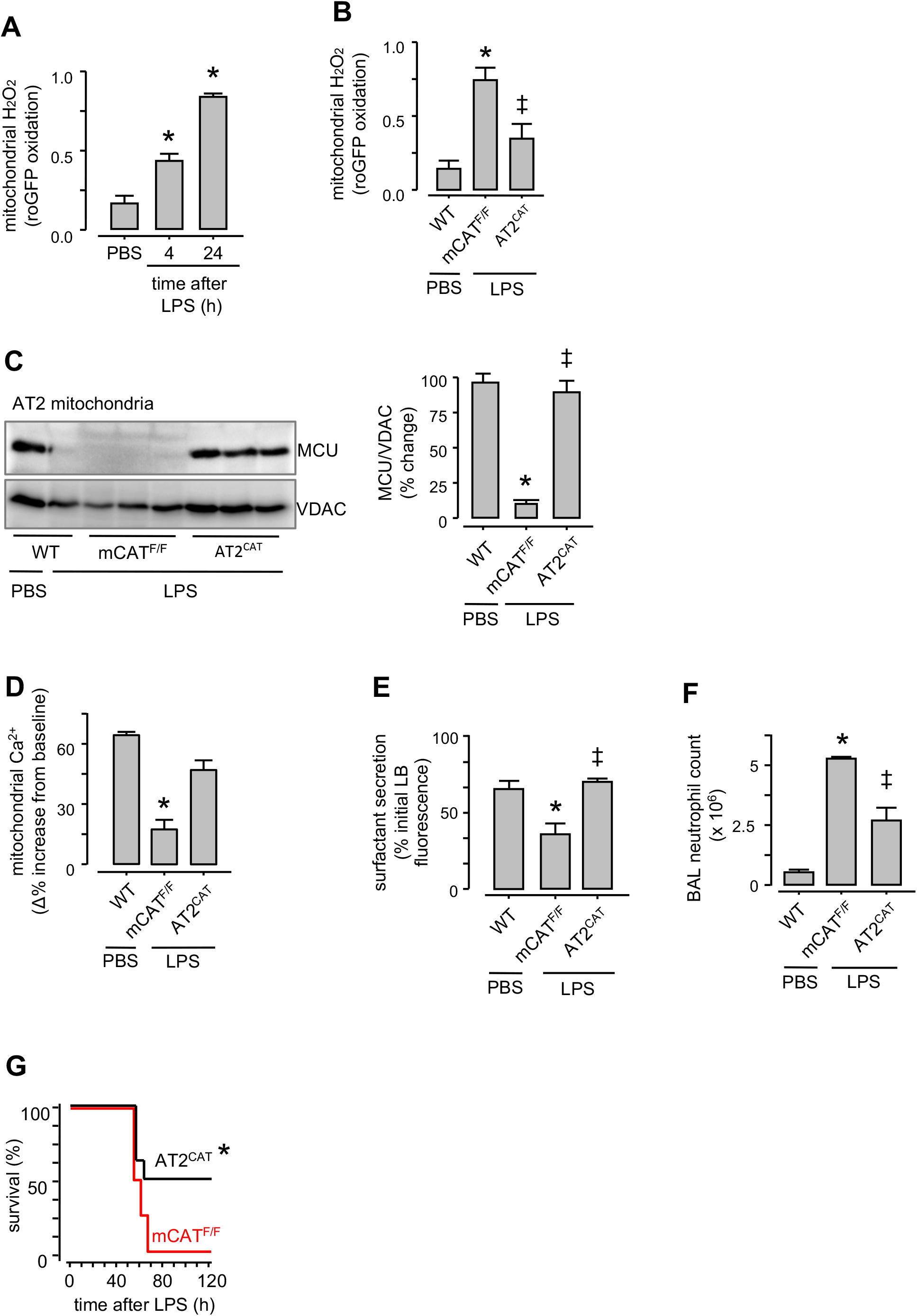
Inhibition of mitochondrial H_2_O_2_ abrogates AT2 MCU depletion. (A and B) Group data are for baseline AT2 mitochondrial H_2_O_2_ production following sublethal LPS instillations. *mCAT^F/F^*, mice floxed for mitochondrial catalase (*mCAT*): *AT2^CAT^*, mice expressing mCAT in AT2. In *B*, determinations were made 24h after indicated instillations. LPS was instilled at a sublethal dose. n=4 lungs for each group, *p<0.05 versus PBS; ‡p<0.05 versus mCAT^F/F^. (C) MCU immunoblots and densitometry of AT2 mitochondria from lungs given intranasal PBS or sublethal LPS. Lungs were excised and AT2 isolated 24h after instillations. n=3 lungs for each group, *p<0.05 versus PBS; ‡p<0.05 versus mCAT^F/F^. (D-F) Group show *in situ* determinations of AT2 MCU function (*D, E*) and lung inflammation (*F*). All determinations were made 24h after intranasal instillations. LPS was instilled at a sublethal dose. n=4 lungs each bar, *p<0.05 versus PBS; ‡p<0.05 versus mCAT^F/F^. (G) Kaplan-Meier plots are for mouse survival after instillations of ARDS-inducing lethal LPS. n=10 mice in each group, *p<0.05 versus mCAT^F/F^. Group data are mean±SEM.

We considered the possibility that mitochondrial H_2_O_2_ might induce mitochondrial fragmentation, and thereby impair mitochondrial Ca^2+^ uptake (Wang et al., 2017). Mitochondrial H_2_O_2_ can activate the fragmentation inducer, dynamin-related protein-1 (Drp1) (Gomes and Scorrano, 2013; Preau et al., 2016). Since mitochondrial distribution has not been determined in intact systems, we imaged AT2 in intact alveoli of PhAM floxed:E2a-Cre mice that globally express the mitochondria-targeted fluorescent protein, Dendra-2 (Luchsinger et al., 2016) (Figures S4A and 4A). AT2 mitochondria aggregated at peri-nuclear poles, co-existing with surfactant-containing lamellar bodies. However, this polarized aggregation was lost by Day 1 after LPS, as fragmented mitochondria distributed throughout the cytosol (Figure 4A,B). However, following intranasal delivery of the Drp1 inhibitor, Mdivi-1 (Manczak et al., 2019), LPS failed to disrupt the polarized mitochondrial distribution (Figure 4A,B). Thus, Drp1 inhibition blocked LPS-induced mitochondrial disaggregation.

**Figure 4.**
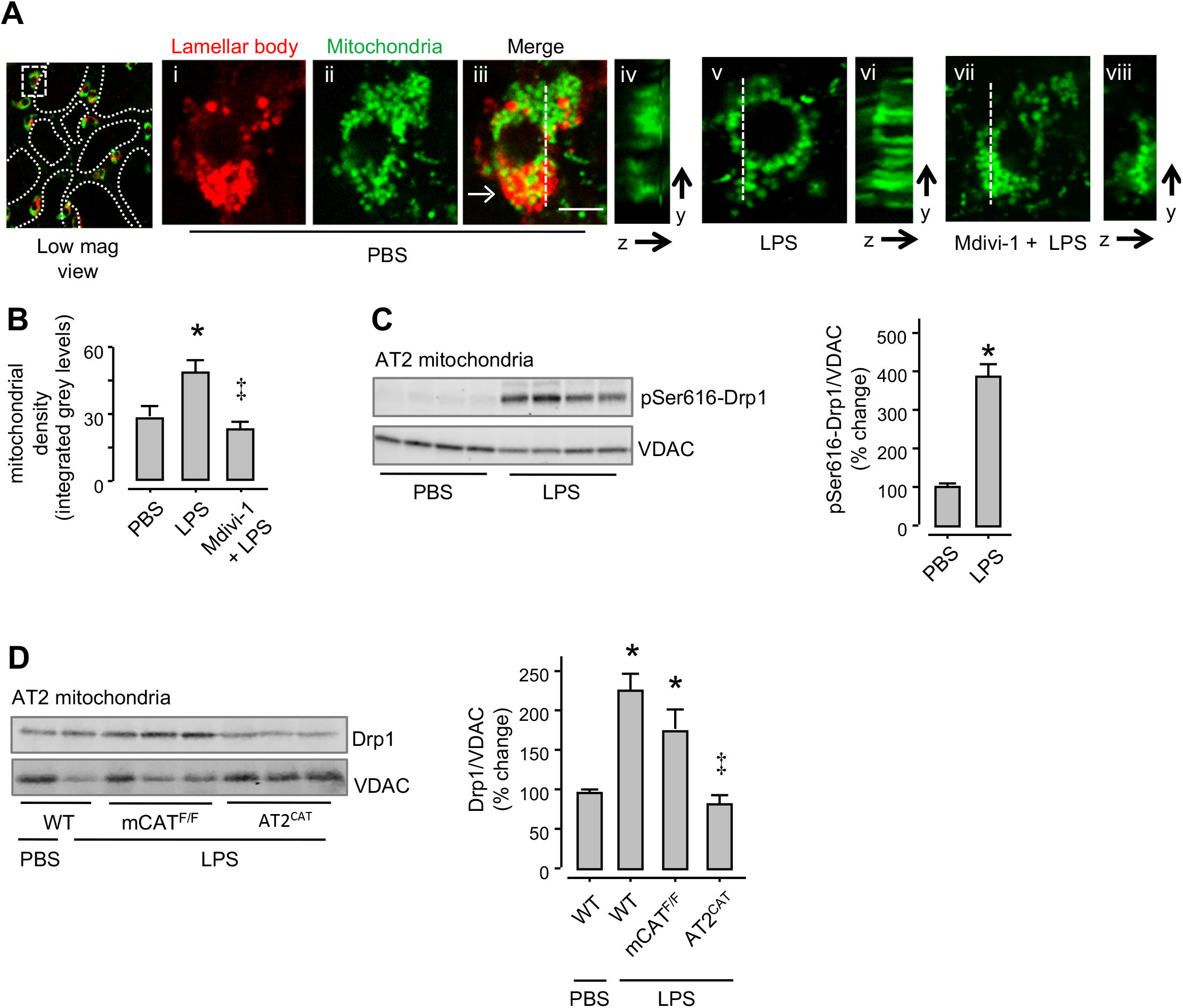
LPS causes mitochondrial fragmentation. (A and B) In mice expressing mitochondria-targeted dendra-2, we detected AT2 in terms of the lamellar body (LB) localizing dye, LTR. In the PBS-treated lung (*alveoli marked by dotted lines*), an AT2 is selected in the low magnification image (*rectangle*). Images *i-iii* show the AT2 at high magnification, displaying polarized mitochondrial aggregation. LBs co-mingle with mitochondria at the indicated site (*arrow*). Images *v* and *vii* display the channel for mitochondrial fluorescence in AT2 in a lung given intranasal instillation of sublethal LPS, and a lung that was given Mdivi-1 prior to the LPS treatment. Images *iv, vi* and *viii* show distribution of mitochondrial fluorescence along the depth axes (*y-z*). Mitochondrial density was quantified in the depth axis at sites of highest mitochondrial aggregation along the selected lines (*dashed lines*). Scale bar, 5μm. n=20 cells from 4 lungs each bar. p<0.05 versus PBS (*), or LPS (‡). (C and D) Phosphorylated Ser616-Drp1 (*C*) and Drp1 (*D*) immunoblots and densitometry are for freshly isolated AT2 mitochondria following indicated treatments. LPS was instilled at a sublethal dose. *Drp1*, dynamin-related protein 1; *pSer616*, phosphorylated serine 616. n=3 lungs each bar, *p<0.05 versus PBS; ‡p<0.05 versus mCAT^F/F^. Group data are mean±SEM.

We confirmed that Drp1 was activated, as revealed by Drp1 phosphorylation at Ser-616 through immunoblots of AT2 mitochondria with a specific antibody (Simula et al., 2018) (Figure 4C). Immunoblots also revealed Drp1 localization to mitochondria in floxed littermate controls, but not in AT2^CAT^ mice (Figure 4D). Thus Drp1 activation was H_2_O_2_ dependent. To evaluate this mechanism further, we induced AT2-specific *Drp1* deletion through post-partum cre recombination in *Drp1*^F/F^:SPC-cre mice (AT2^*Drp1*-^). We confirmed the deletion by immunoblot of isolated AT2 (Figure S4B). In AT2^*Drp*-^ mice, there was no LPS-induced MCU loss (Figure 5A). The MCU loss was also inhibited in mice with AT2 expression of a Drp1 mutant that lacks GTPase activity (Cereghetti et al., 2010; Smirnova et al., 2001) (Figure 5B), or after pre-treatment with Mdivi-1 (Figure S4C). These findings indicated that Drp1activation caused the MCU loss.

**Figure 5.**
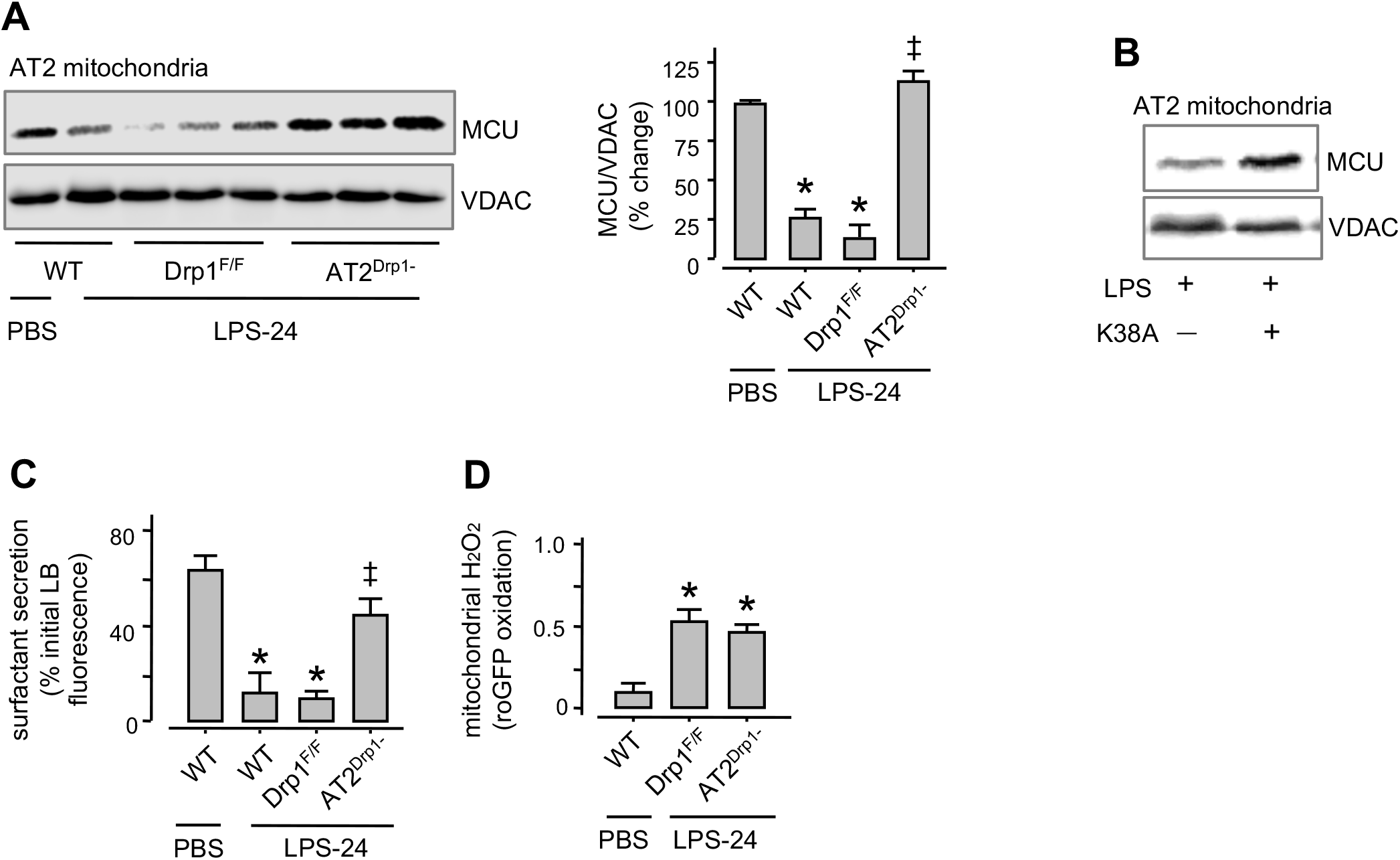
LPS causes H_2_O_2_ induced Drp1 activation. (A) MCU Immunoblots and densitometry are shown for AT2 mitochondria derived from lungs of mice given the indicated treatments. LPS was instilled at a sublethal dose. *Drp1^F/F^*, floxed mice for Drp1; *AT2^Drp1-^*, mice lacking Drp1 in AT2. n= 3 lungs for each bar. Lungs were excised and AT2 isolated 24h after intranasal instillations. *p<0.05 versus corresponding PBS; ‡p<0.05 versus Drp1^F/F^. (B) MCU immunoblots in AT2 mitochondria derived from mice expressing an empty vector or a kinase-dead Drp1 (*K38A*). All animals were given intranasal LPS at a sublethal dose and lungs excised 24h after instillation. n= 3 lungs for each group. (C and D) Group data show respectively, surfactant secretion (*C*) and mitochondrial H_2_O_2_ (*D*) following indicated treatments. All determinations were made 24h after instillations. n= 4 lungs for each bar. *p<0.05 versus corresponding PBS; ‡p<0.05 versus Drp1^F/F^. Group data are mean±SEM.

Since MCU loss inhibited surfactant secretion, we evaluated the secretion in LPS-treated AT2^*Drp1*-^ mice. Since MCU was not lost in these mice, surfactant secretion was robust (Figure 5C). Since H_2_O_2_ production was upsteam of the Drp1 activation, the LPS-induced H_2_O_2_ increase was present in AT2^*Drp1*-^ mice (Figure 5D). Thus, we show that LPS exposure induced a chain of events in AT2 mitochondria, in which H_2_O_2_-induced Drp1 activation caused mitochondrial fragmentation and loss of MCU. The resulting impairment of surfactant secretion likely contributed to the dysfunctional lung fluid balance underlying ARDS.

### Factors contributing to mitochondrial responses in AT2

In LPS-treated lungs episodic AT2 cCa^2+^ increases occur because of Ca^2+^ conduction through connexin 43 (Cx43)-containing gap junctions (GJs) in the alveolar epithelium (AE) (Ashino et al., 2000; Ichimura et al., 2006). Cx43^F/F^:SPC-cre mice lack Cx43 in the AT2 (AT2^*Cx43*-^) (Figure S5A). As opposed to floxed littermate controls, in AT2^*Cx43*-^ mice LPS failed to decrease MCU (Figure 6A) and mitochondrial buffering was robust (Figure 6B). These findings indicated that the presence of AE GJs supported the MCU loss.

**Figure 6.**
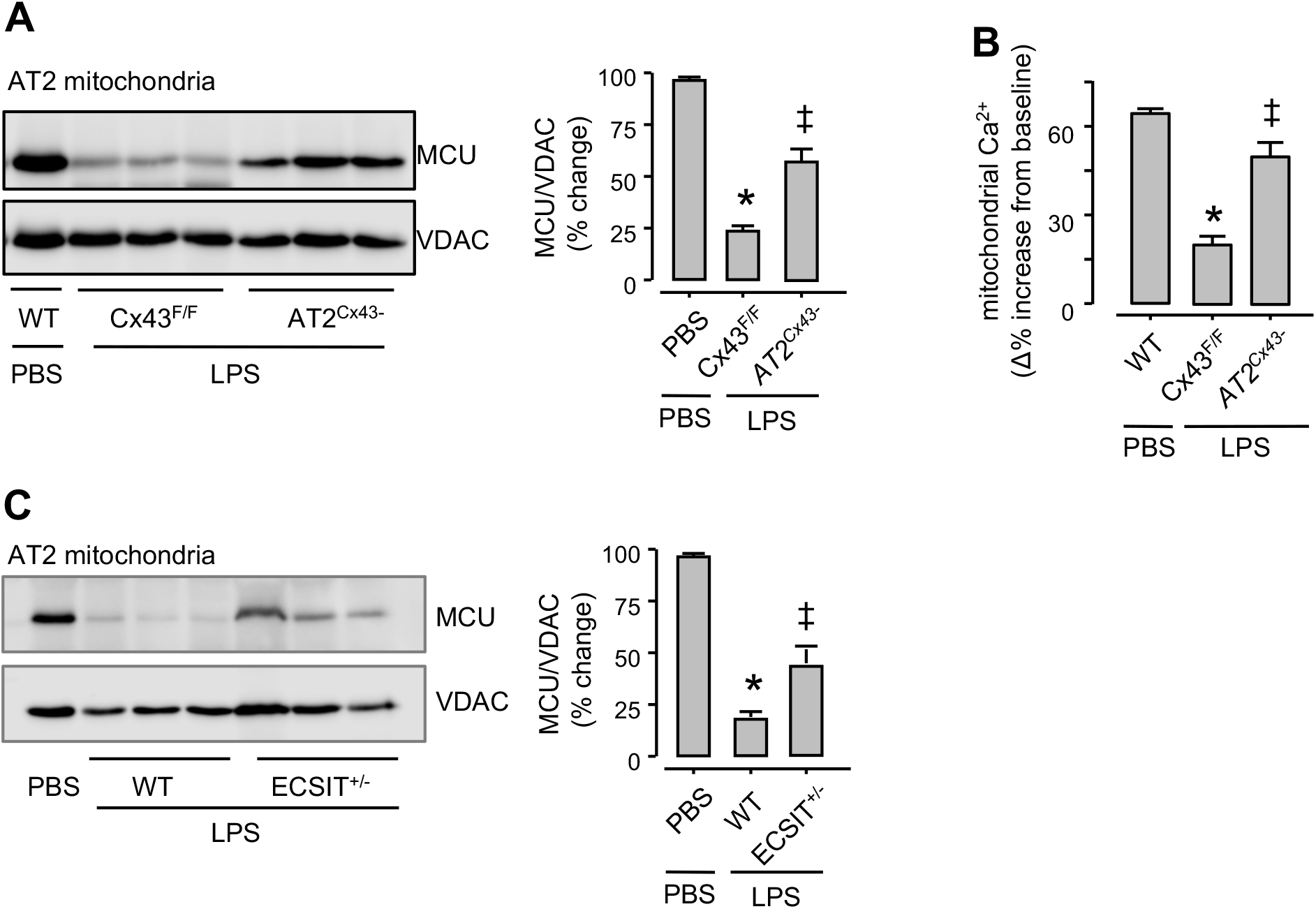
Mechanisms of mitochondrial H_2_O_2_ generation. (A) Immunoblots in AT2 mitochondria (*left*) and densitometric quantification of immunoblots (*right*) show MCU expression 24h after indicated treatments. LPS was instilled at a sublethal dose. *Cx43*, connexin 43; *Cx43^F/F^*, mice floxed for *Cx43*; *AT2^Cx43-^*, mice lacking *Cx43* in AT2. n=4 lungs each bar, *p<0.05 versus PBS; ‡p<0.05 versus Cx43^F/F^. (B) Determinations of mitochondrial Ca^2+^ responses to alveolar stretch 24h after indicated intranasal instillations. LPS was instilled at a sublethal dose. n=4 lungs each bar, *p<0.05 versus PBS; ‡p<0.05 versus Cx43^F/F^. (C) Immunoblot (left) and densitometric determinations (right) from AT2 mitochondria. AT2 were isolated 24h after instillations of either PBS or sublethal LPS. *ECSIT*, evolutionarily conserved signaling intermediate in Toll pathway; *ECSIT^+/-^*, heterozygous knockout mice for ECSIT. n=3 lungs each bar, *p<0.05 versus PBS; ‡p<0.05 versus WT.

It is proposed that LPS-induced activation of Toll-like receptor (TLR4) translocates the TLR-signaling adaptor molecule, TRAF6 to mitochondria (Kopp et al., 1999). TRAF6 forms a complex with Complex I assembly factor, ECSIT (evolutionarily conserved signaling intermediate in Toll pathways), augmenting mitochondrial H_2_O_2_ formation (West et al., 2011). We tested this possibility in mice with heterozygous knockout of ECSIT (ECSIT^+/-^) in which ECSIT expression is 50% of wild type (West et al., 2011). In ECSIT^+/-^ mice, the LPS-induced MCU loss was markedly mitigated (Figure 6C). Taken together, our findings indicate that Cx43-containing AT2 GJs and mitochondrial ECSIT contributed to the MCU loss.

### Rescue of AT2 MCU protects against LPS-induced mortality

To further evaluate the MCU role, we replenished *MCU* in in mice globally lacking the MCU (*MCU^-/-^*) through intranasal instillation of a lentiviral vector for MCU transduction (Pan et al., 2013). The viral transduction increased MCU expression in AT2 (Figure S6A,B). Although LPS-induced mortality in *MCU^-/-^* mice was two times above control (Figure 7A), *MCU* transduction markedly abrogated the mortality (Figure 7A), affirming the protective effect of the MCU replenishment.

**Figure 7.**
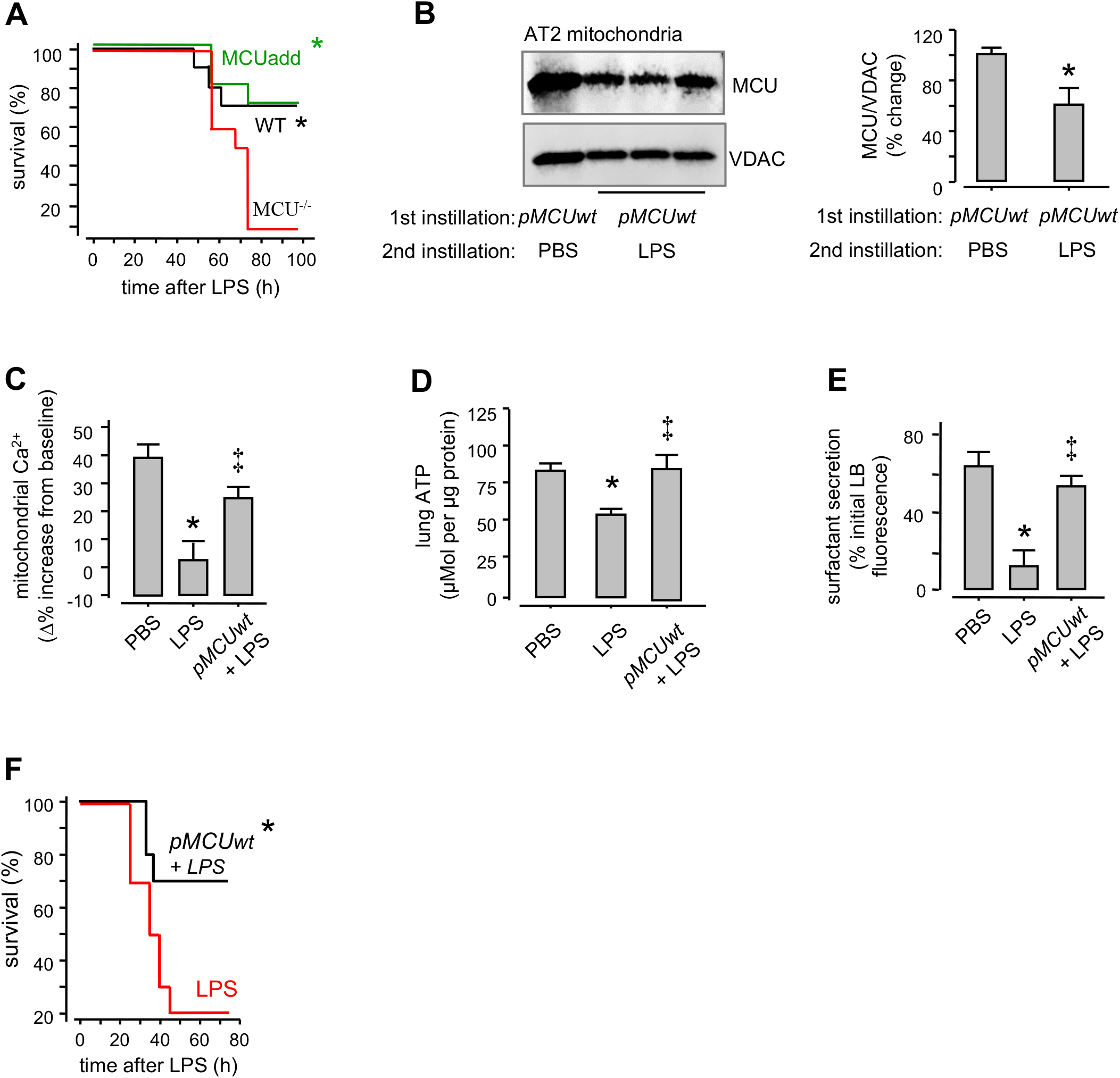
Effects of wild-type *MCU* overexpression in alveolar epithelium. (A) Kaplan-Meier plots for mouse survival in indicated groups after instillations of moderate-dose LPS in wild-type (*WT*), MCUnull (*MCU^-/-^*), and MCU^-/-^ mice following add-back of MCU (*MCUadd*). To add-back MCU, we intranasally instilled a lentiviral vector encoding for the wild-type MCU in MCU^-/-^ mice. LPS was instilled at a lethal dose 14d after lentivirus instillation. n=15 mice in each group, *p<0.05 versus MCU^-/-^. (B) Immunoblots and densitometric quantifications of AT2 mitochondria from lungs given the indicated intranasal instillations. Liposome-complexed *pMCUwt* plasmid (plasmid encoding for full length MCU) was given by intranasal instillation. Instillation sequences were: PBS followed 8h later by *pMCUwt* for the control group, and*pMCUwt* followed 24h later by instillation of sublethal LPS for the treated group. Lungs were removed for analyses 48h after the first instillations. n=3 lungs for each bar, *p<0.05 versus PBS. (C - E) Quantifications of mitochondrial Ca^2+^ (*C*), lung ATP (*D*) and surfactant secretion (*E*) were obtained 24h after intranasal instillations of PBS or sublethal LPS in wild-type or in *pMCUwt* expressing mice. *pMCUwt*-complexed liposomes were intranasally instilled 24h before instillations of sublethal LPS. Lung ATP was determined by a colorimetric method. n=4 lungs per group, *p<0.05 versus PBS and ‡p<0.05 versus LPS. (F) Kaplan-Meier plots are for mouse survival in Swiss Webster mice following instillations of lethal dose LPS followed 24h later with PBS (*LPS*) or *pMCUwt* followed 24h later with LPS (*pMCUwt*). n=10 mice in each group, *p<0.05 versus LPS. Group data are mean±SEM.

To determine the effects of *MCU* overexpression in wild-type mice, we instilled liposome-complexed *MCU* plasmid, then confirmed that 48h after transfection, AT2 MCU expression was two times above baseline (Figure S6C,D). Subsequently, as expected from our findings above, LPS decreased the MCU expression in LPS-24 lungs (Figure 7B). However, despite this decrease, the expression remained sufficiently high (60% of baseline) that buffering was retained (Figure 7B,C). Although LPS caused the expected decrease of ATP in controls (Islam et al., 2012), *MCU*-transfection protected ATP production (Figure 7D), surfactant secretion (Figure 7E), and survival (Figure 7F). Together, these studies indicated that in LPS-induced ARDS, MCU overexpression ensured adequate protection of AT2 MCU function, thereby protecting survival.

As a possible therapeutic approach for enriching AT2 MCU in ARDS, we considered a mitochondrial transfer strategy. Mitochondrial transfer from bone marrow-derived mesenchymal stromal cells (BMSCs) to the alveolar epithelium protects against ARDS (Islam et al., 2012). However, the protective mechanisms are not understood. To determine the role of the AT2 MCU in this protection, we transfected BMSCs with plasmids encoding GFP-tagged, wild-type MCU (*MCUwt*) (Figure S7A), or a mutant MCU (*MCUmt*) that inhibits mitochondrial Ca^2+^ entry (Raffaello et al., 2013) (Figure S7B). We intranasally instilled BMSCs expressing the *MCU* plasmids 4h after instilling LPS. In LPS-24 lungs of these mice, we confirmed that BMSC mitochondria were transferred to AT2 (Figure S7C,D), and that the transfer increased MCU expression in AT2 (Figure 8A). Overexpression of *MCUwt* protected MCU function and abrogated inflammation, increasing survival (Figure 8B-E). Overexpression of *MCUmt* failed to achieve the protections. Taking our findings together, we conclude that primary lack of the AT2 MCU was sufficient to tilt the inflammatory outcome towards ARDS, and that rescue of MCU function by mitochondrial transfer protected inflammation resolution and survival.

**Figure 8.**
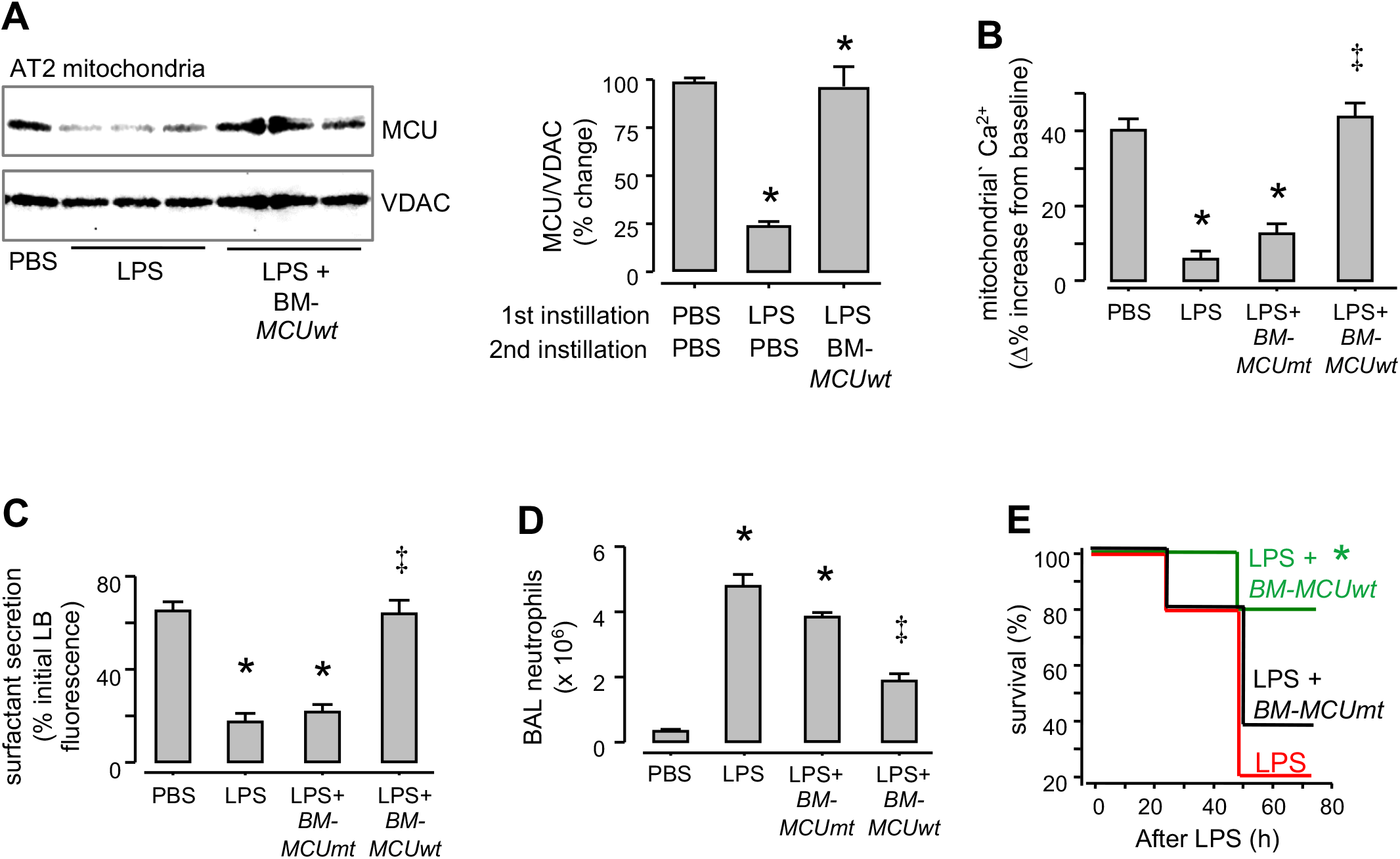
BMSC expression of *pMCUwt* protects against LPS injury. (A) MCU immunoblots in AT2 mitochondria derived from mice given indicated intranasal instillations. BMSCs expressing the wild-type MCU plasmid (*BM-MCUwt*) were intranasally instilled 4h after sublethal LPS instillations. Lungs were excised and AT2 isolated 24h after LPS instillations. Bars show densitometry. n= 3 lungs for each group, *P<0.05 versus bar on left. (B, C) Group data are AT2 mitochondrial Ca^2+^ (*B*) and surfactant secretion (*C*, loss of LTR fluorescence) responses following alveolar stretch. Sequence of intranasal instillations is indicated. All determinations were carried out 24h after the first instillation. *BM-MCUmt*, BMSCs expressing a mutant MCU. n=4 lungs each bar, *p<0.05 versus PBS and ‡p<0.05 versus LPS. (D) Bars quantify the alveolar inflammatory response to indicated instillations as indicated by neutrophil counts in the bronchioalveolar lavage (BAL) obtained 24h after the first instillation. *p<0.05 versus PBS and ‡p<0.05 versus LPS. (E) Kaplan-Meier plots for mouse survival after instillations of LPS at a lethal dose in Swiss Webster mice. The groups are: LPS alone (*red*), LPS followed 4h later with instillation of BM-MCUwt (*green*) and BM-MCUmt (*black*). n=5 mice in each group, *p<0.05 versus LPS alone. Group data are mean±SEM.

## DISCUSSION

We report here our finding that viability of MCU function in the alveolar epithelium determines ARDS severity. Direct evidence of this was obtained by genetic deletion studies, showing that tissue-specific knockout of the MCU in the alveolar epithelium markedly exacerbated LPS-induced mortality. The MCU loss in the alveolar epithelium occurred as part of the innate immune response, since exposing mice to a sub-lethal LPS dose, or to *P, aeruginosa* progressively decreased the MCU expression. Importantly, survival depended on spontaneous or experimental replenishment of the MCU expression. MCU reinstatement restored mitochondrial buffering in AT2, re-establishing surfactant secretion, mitigating inflammation and abrogating mortality. Thus, alveolar MCU was revealed to be critical for survival during immune stress.

Treating mice with a sub-lethal LPS dose enabled recognition of mitochondrial responses in a time-dependent manner. The time course of the MCU expression was unchanged from baseline for about 8h post-LPS. Subsequently however, the expression progressively decreased until it was undetectable by Day 1. These time courses tallied with indices of MCU function, which were intact early after LPS exposure, despite the onset of a robust immune response. Notably however, both MCU expression and MCU function failed by Day 1. Concomitantly, mitochondrial H_2_O_2_ production progressively increased. Expression of AT2-specific catalase blocked the MCU loss, implicating mitochondrial H_2_O_2_ as a determinant of MCU expression.

Increase of mitochondrial H_2_O_2_ in AT2 can occur as a result of Complex 1 instability induced by binding of TRAF6 to ECSIT (Kopp et al., 1999; West et al., 2011), or by Ca^2+^ communication across Cx43-containing GJs in the AE (Ashino et al., 2000; Ichimura et al., 2003). We affirmed these mechanisms, in that heterozygous ECSIT knockout, or AT2-specific Cx43 knockout each markedly abrogated the LPS-induced MCU loss. These mechanisms may have operated in tandem, the cCa^2+^-induced effect operating in the early phase of the immune response when mitochondrial buffering was present. In the later phase, as buffering failed, the ECSIT-induced effect may have dominated. Together, these mechanisms may have established the MCU-degrading, sustained H_2_O_2_ increase.

The H_2_O_2_ production activated Drp1, an effect that was inhibited by expression of mitochondrial catalase. Importantly, MCU loss was blocked in mice lacking *Drp1* in AT2. Taking these findings together, we conclude that H_2_O_2_-activation of Drp1 was the critical mechanism underlying the LPS-induced MCU loss. We speculate that during immune challenge, the cell decreases MCU expression as negative feedback against further mCa^2+^-induced oxidant production.

Drp1 activation, resulting from exposure to viruses, bacteria and other inflammogens (Cui et al., 2018; Jain et al., 2011; Kim et al., 2014; Park et al., 2018) is implicated in the mitochondrial fragmentation that precedes mitophagy, a process by which the cell eliminates dysfunctional mitochondria (Gomes and Scorrano, 2013). Here, LPS caused mitochondrial fragmentation as indicated by a loss of their polarized aggregated formations, and to absence of fusion-fission activity. However, contents of several proteins of the mitochondrial inner and outer membranes, as well as the mitochondrial membrane potential, a measure of mitochondrial fitness, were unchanged. Further, parkin localization to mitochondria, a marker of mitophagy initiation (Cereghetti et al., 2010; Matsuda et al., 2010), was undetectable. Hence, we interpret that the fragmentation did not cause loss of AT2 mitochondria, and that the LPS-induced MCU loss was not due to Drp1-activated mitophagy of AT2. In unstressed lungs, AT2 mitochondria clustered with LBs, suggesting that the close LB-mitochondria proximity facilitated transfer of mitochondrial ATP to LBs to activate surfactant secretion. Diminished ATP transfer following mitochondrial disaggregation might account for inhibition of LPS-induced inhibition of surfactant secretion.

In conclusion, our findings attest to the critical role played by the AT2 MCU in protecting mitochondrial buffering for the maintenance of alveolar stability. Evidently, the unprotected cCa^2+^ increases that occurred in the buffering-compromised alveolar epithelium, accounted for two major injury mechanisms. First, there was compromise of the epithelial fluid barrier, as evident in the increases of the extravascular lung water content. The barrier loss might be have been due cCa^2+^-induced actin depolymerization, (Hough et al., 2019). Second, MCU deficiency inhibited mCa^2+^-induced ATP increases required for surfactant secretion. These injury mechanisms likely induced the pulmonary edema that exacerbated mortality. We propose that restoration of AT2 MCU function by mitochondrial replenishment therapy might provide a strategy for the prevention and reversal of ARDS.

## ACKNOWLEDGMENTS

Michelle Wei assisted with genotyping. The PhAM floxed:E2a-Cre mice were a gift of Dr. Hans Snoeck (Columbia University). Purified human SPB was a gift of Dr. Timothy Weaver (University of Cincinnati). This work was supported by NIH grants HL36024, HL57556, and HL122730 to JB, and a Parker B. Francis Fellowship and an American Heart Association Grant-in-Aid to MNI. Some studies were carried out in the Columbia Center for Translational Immunology Flow Cytometry Core, supported in part by NIH award S10OD020056.

## AUTHOR CONTRIBUTIONS

M.N.I. planned, carried out and analyzed all experiments. G.A.G. contributed to alveolar type 2 cell isolation and imaging experiments. L. L. and E.M. contributed to alveolar stretch experiments. M.A. and E.O-A. performed mitochondrial complex activity determinations. S.D. carried out immunoblotting. S.B. contributed to the plan. J.B. designed the overall project. All authors contributed to the writing.

## DECLARATION OF INTERESTS

The authors declare no competing interests.

## STAR METHODS

Detailed methods are provided in the online version of this paper and include the following:

- KEY RESOURCES TABLE
- CONTACT FOR REAGENT AND RESOURCE SHARING
- EXPERIMENTAL MODEL AND SUBJECT DETAILS

○ Animal methods
○ ARDS methods
○ Bone marrow-derived mesenchymal stromal cell (BMSC) isolation, purification and culture
- METHOD DETAILS

○ Isolated, blood-perfused lungs
○ Alveolar microinfusion and imaging
○ Alveolar immunofluorescence
○ AT2 isolation by flow cytometry
○ Mitochondria isolation
○ Immunoblotting/immunoprecipitation
○ RNA determination
○ siRNA and plasmid transfection in mice
○ Lentivirus cloning and production
○ Channelrhodopsin activation
○ Mitochondrial complex activity assay
○ Extravascular lung water
○ Lung ATP determination
○ BAL neutrophil count
- QUANTIFICATION AND STATISTICAL ANALYSIS

## STAR METHODS

### KEY RESOURCE TABLE

**Table 1.**
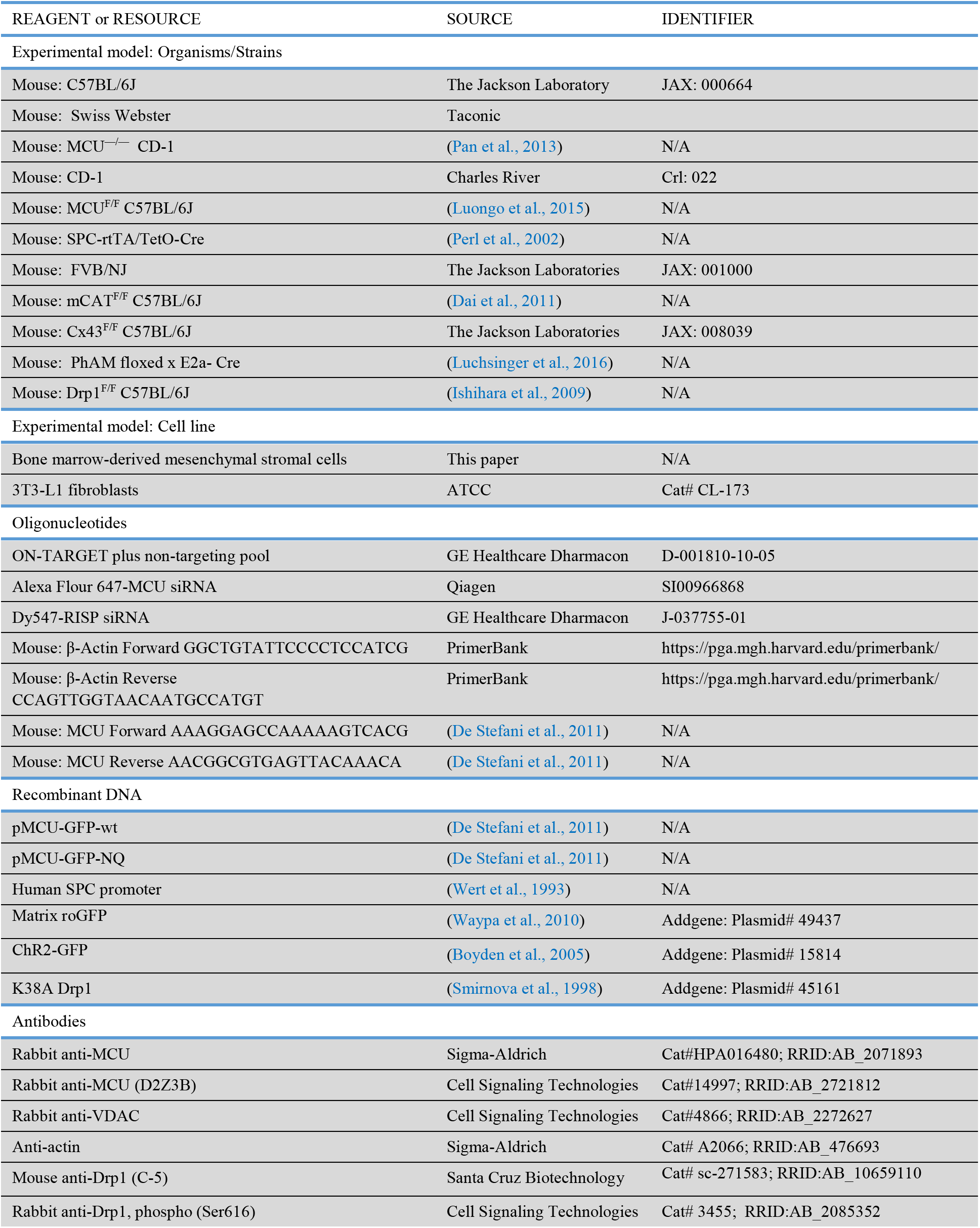
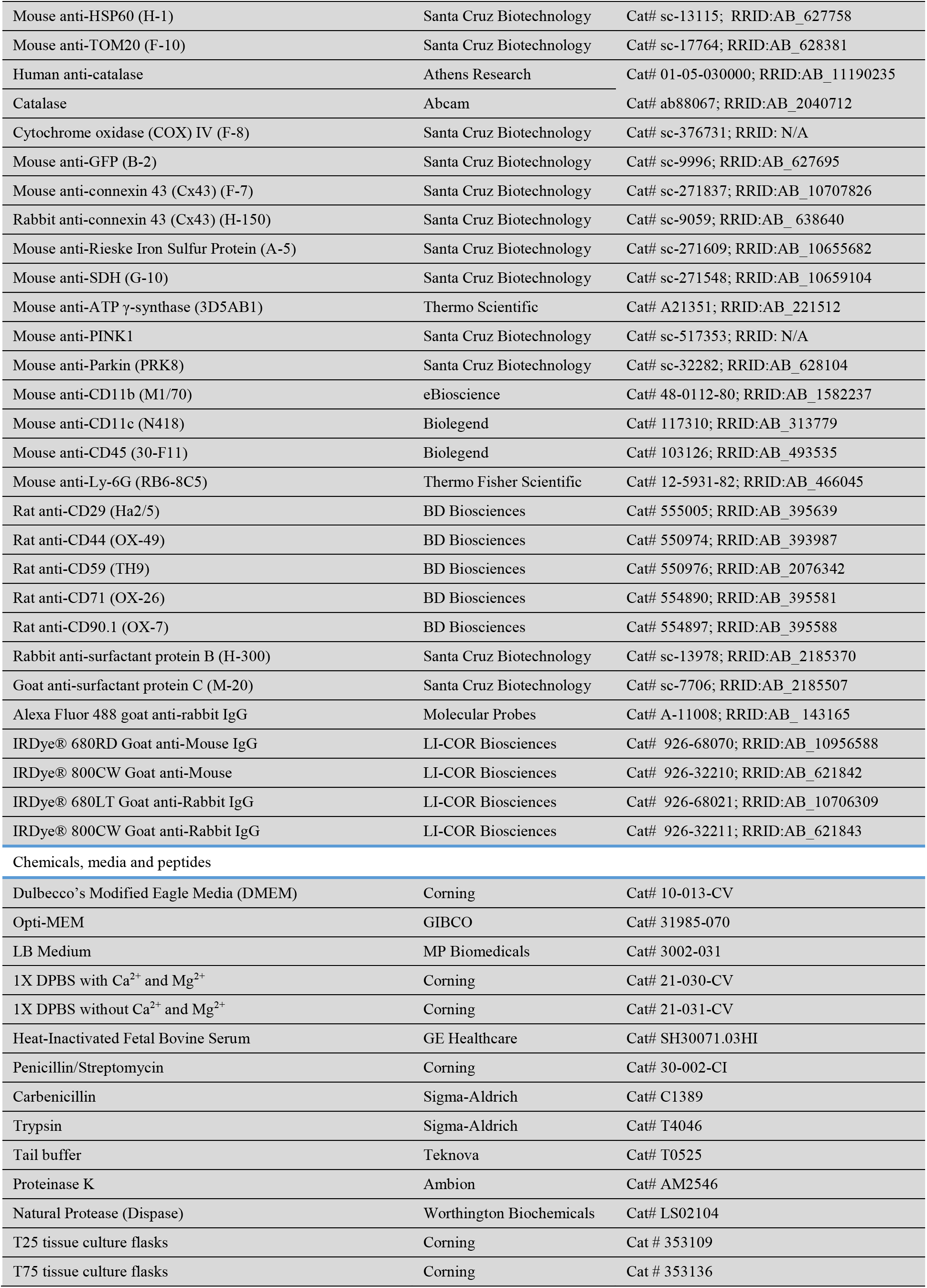
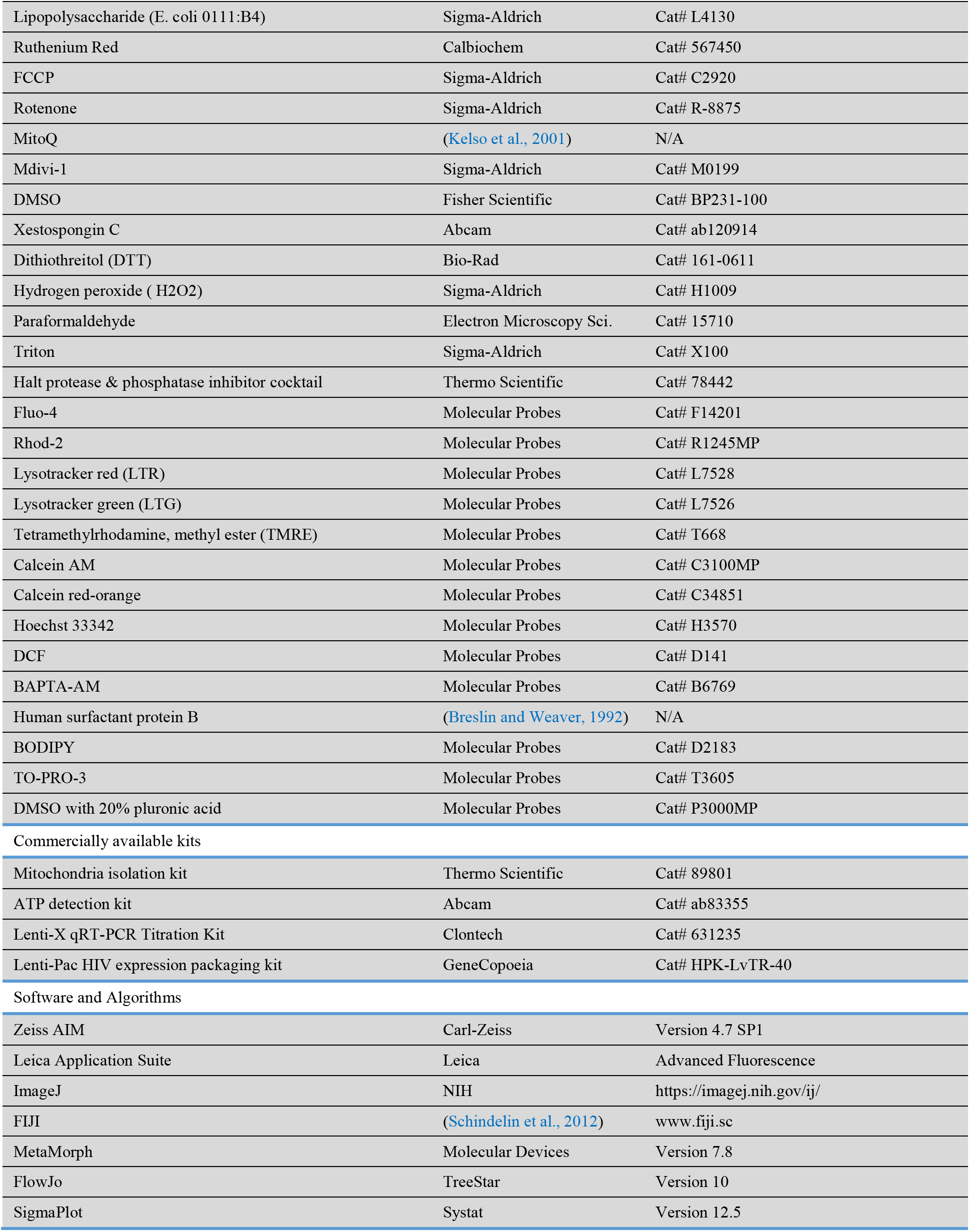

### CONTACT FOR REAGENTS AND RESOURCE SHARING

Further information and requests for resources and reagents should be directed to and will be fulfilled by the Lead Contact, Dr. Jahar Bhattacharya. MTAs were obtained for the transfer of MCU^F/F^, MCU^-/-^, mCAT^F/F^, Drp1^F/F^ and SPC-Cre mice.

### EXPERIMENTAL MODEL AND SUBJECT DETAILS

#### Animal methods

Animal procedures were approved by the Institutional Animal Care and Use Committ of Vagelos College of Physicians and Surgeons at Columbia University. All animals were cared for according to the NIH guidelines for the care and use of laboratory animals. Mice were socially housed under a 12h light/dark cycle with *ad libitum* access to water and food. Age and sex matched animals wei randomly assigned to experimental groups by investigators. Investigators were blinded for the all surviv and BAL leukocyte count studies.

#### Bacterial culture

We cultured single colonies of *Pseudomonas aeruginosa* (strain K, provided by A. Prince, Columbia University, New York, NY, USA), overnight in 4 ml of Luria-Bertani (LB) medium a 37°C and 250 rpm (Innova42, New Brunswick Scientific). *P. aeruginosa* were selected with Carbenicillin (300 μg/ml). On the day of the experiment, overnight cultures were diluted 1:100 in fresh LB and grown in a shaking incubator to optical density (OD) 0.5 at 600 nm (SPECTRAmax Plus, Molecular Devices).

#### In vivo bacterial instillation

For intranasal instillations in mice, 1 ml bacterial culture was centrifuged and resuspended in 10 ml sterile PBS. In anesthetized (ketamine-100 mg/kg and xylazine 5 mg/kg, i.p.) animals, we airway instilled 50 μl suspension to deliver “high” inoculum (1 x 10^6^ CFU) per mouse. For “low” inoculum instillation, we diluted (x10) the “high” inoculum suspension, then airway instilled 50 μ suspension to deliver 1 x 10^5^ CFU per mouse.

#### Adult Respiratory Distress Syndrome (ARDS) methods

ARDS was induced in anesthetized (ketamine-100 mg/kg and xylazine 5 mg/kg, i.p.) animals by airway instillation of LPS (*E.coli* 0111.B4) in sterile PBS. Control animals were instilled an equal volume of sterile PBS. Since multiple mouse strains were used, and since the lung injury-causing LPS dose varies between mouse strains, we gave different LPS doses to simulate sublethal, moderate and lethal ARDS-inducing that caused mortality of 0, 30 and 80% respectively, in control groups (Table 2). The strains are indicated.

**Table 2.**
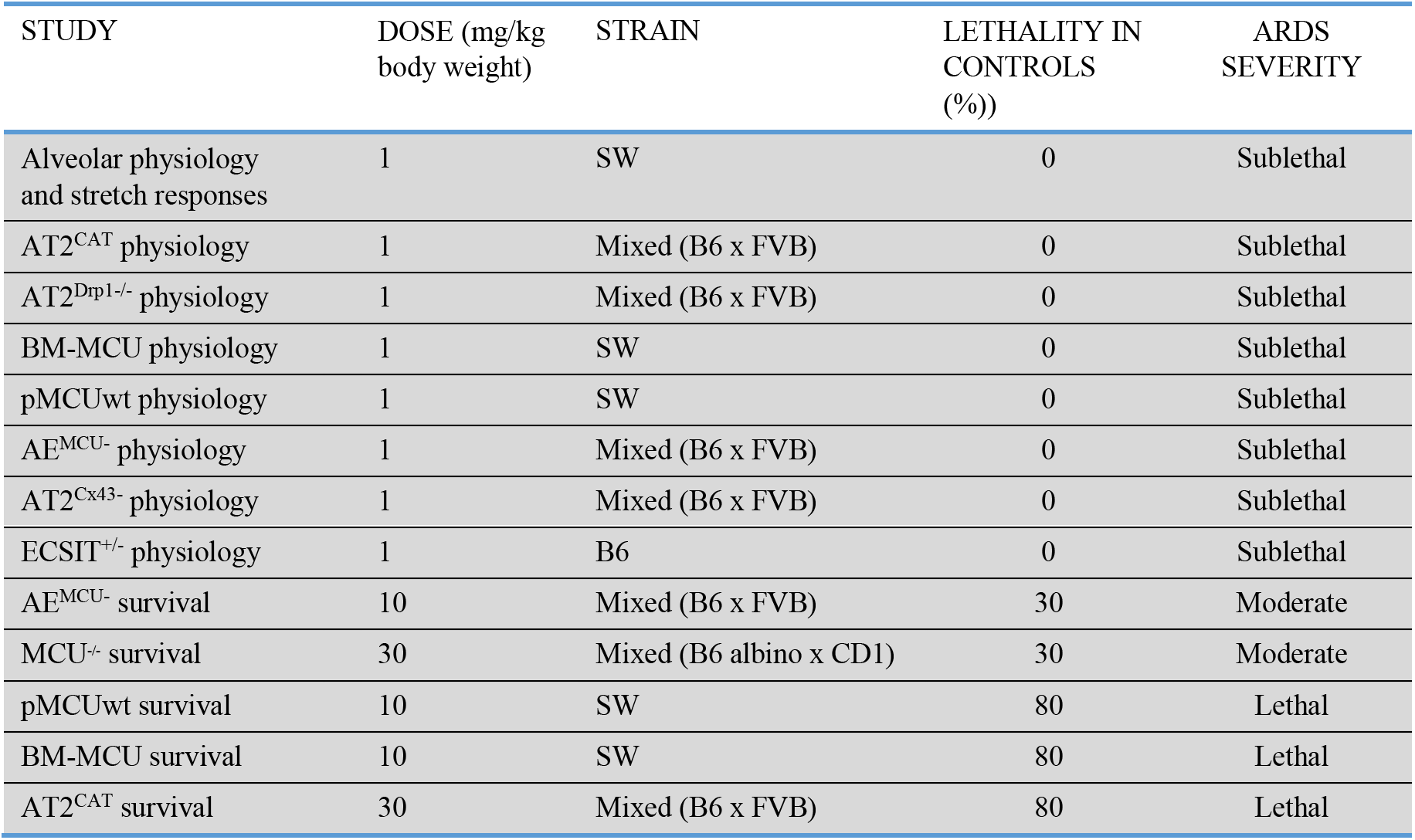

#### Bone marrow-derived stromal cell (BMSC) isolation, purification and culture

We isolated mouse BMSCs according to our reported methods (Islam et al., 2012). Briefly, in anesthetized animals, bone lumens of excised femurs and tibia were infused with MSC growth medium to recover marrow. Three days after marrow plating on tissue culture flasks, the supernatant was rejected and adherent BMSCs were incubated (37°C) under subconfluent conditions to prevent cell differentiation. BMSCs from passages 5 to 12 were used in this study. In accordance with established criteria (Dominici et al., 2006), BMSCs were characterized as we reported (Islam et al., 2012). BMSCs were cultured in 5% CO_2_ at 37°C in DMEM containing 10% FBS and 1% of an antibiotic mixture.

### METHOD DETAILS

#### Isolated, blood-perfused lungs

Using our reported methods (Islam et al., 2012; Westphalen et al., 2014), lungs were excised from anesthetized mice (i.p., Ketamine 80-100 mg/kg, Xylazine 1-5 mg/kg), then perfused with autologous blood through cannulas in the pulmonary artery and left atrium. The blood was diluted in 4% dextran (70 kDa), 1% fetal bovine serum and buffer (150 mmol/l Na^+^, 5 mmol/l K^+^, 1.0 mmol/l Ca^2+^, 1 mmol/l Mg^2+^, and 20 mmol/l HEPES at pH 7.4). The perfusion flow rate was 0.5 ml/min at 37°C at osmolarity of 300 mosM (Fiske Micro-Osmometer, Fiske^®^ Associates, Norwood, MA), and hematocrit 10%. The lung was inflated through a tracheal cannula with a gas mixture (30% O_2_, 6% CO_2_, balance N_2_). Vascular (artery and vein at 10 and 3 cmH_2_O) and airway (5 cmH_2_O) pressures were held constant during microscopy.

#### Alveolar microinfusion and imaging

To load the alveolar epithelium with fluorescent dyes or antibodies, we micropunctured single alveoli with glass micropipettes (tip diameter 3-5 μm) and microinfused ~10 neighboring alveoli (Lindert et al., 2007; Wang et al., 2001). After the microinfusions, the free liquid in the alveolar lumen drained in seconds re-establishing air-filled alveoli (Wang et al., 2001). This rapid clearance indicates that the micropuncture does not rupture the alveolar wall, and that the micropunctured membrane rapidly reseals as reported for other cells (Steinhardt et al., 1994). Nevertheless, we selected non-micropunctured alveoli for imaging. To determine whether the fluorescence detected in the epithelium was intracellular or extracellular, in some experiments we microinfused alveoli for 10 minutes with trypan blue (0.01% w/v), which eliminates extracellular fluorescence (Rowlands et al., 2011). In all experiments in which we infused multiple dyes, we confirmed absence of bleed-through between fluorescence emission channels. We imaged intact alveoli of live lungs with laser scanning microscopy (LSM 510 META, Zeiss and TCS SP8, Leica)

#### Alveolar immunofluorescence

We used our reported methods to detect intracellular immunofluorescence in live alveoli (Islam et al., 2014; Westphalen et al., 2014). Briefly, we gave alveoli successive 20-minute microinfusions of 4% paraformaldehyde and 0.1% triton X-100. Then, we microinfused fluorescence-conjugated antibodies (40 ng/ml) for 10 minutes. In some experiments, microinfusion of antibodies were followed by microinfusion of fluorescence-conjugated secondary antibodies (40 ng/ml). To washout unbound fluorescence, we microinfused buffer for 10 minutes and commenced imaging after a further 10 minutes.

#### AT2 isolation by flow cytometry

We isolated AT2 by our reported methods (Islam et al., 2012). Briefly, isolated lungs were buffer perfused through vascular cannulas to clear blood, then we exposed the lungs to intratracheal dispase (0.2 U/ml, 2ml, 45 minutes) at room temperature. The tissue was suspended in PBS and sieved and the sieved sample was then centrifuged (300g x 5 min). The pellet was resuspended and incubated together with the AT2 localizing dye, lysotracker red (LTR) (Ashino et al., 2000), the nuclear dye, Hoechst 33342 and fluorescence Allophycocyanin (APC)-conjugated Ab against the leukocyte antigen, CD45. The suspension was then subjected to cell sorting (Influx cell sorter) to recover AT2.

#### Mitochondria isolation

We used a commercially available isolation kit (Rowlands et al., 2011) (Thermo Scientific). Briefly, we homogenized lungs and AT2 (Tissue Tearor; Biospec Products, Bartlesville, OK), on ice with buffers provided in the kit. The buffers were supplemented with protease and phosphatase inhibitors. We centrifuged the homogenate (800g x 10 min) and collected the supernatants, which were further centrifuged at 17000g for 15 min at 4°C. The pellets and the supernatants contained respectively, the mitochondrial and the cytosolic fractions. Purity of cell fractionation was determined by immunoblotting for voltage-dependent anion channel (VDAC) in mitochondrial and cytosolic fractions.

#### Immunoblotting/Immunoprecipitation

For immunoprecipitation and immunoblotting, mitochondrial fractions were lysed in 2% SDS. For immunoprecipitation, lysates containing 2 mg total protein were precleared with appropriate control IgG for 30 minutes at 4°C with 20 μl of prewashed protein A/G and processed as previously described (Islam et al., 2012). Equal amounts (100 μg) of protein from lysates were separated by SDS-PAGE, electro-transferred onto nitrocellulose membrane overnight at 4°C and blocked in Starting Block Blocking Buffer (Pierce) for 1h then subjected to immunoblotting. Densitometry was performed using ImageJ software.

#### RNA determination

RNA was extracted from isolated AT2 using the RNeasy Micro Kit. The purity of the RNA was assessed by absorbance at 260 and 280 nm using a Thermo Scientific NanoDrop spectrophotometer. Using RNA with a 260/280 ratio of > 1.8, cDNA was synthesized using oligo (dT) and Superscript II. Quantitative RT-PCR was performed using a 7500 Real-Time PCR system (Applied Biosystems) and SYBR Green Master Mix.

#### siRNA and plasmid transfection in mice

We purchased siRNA against MCU and RISP. For knockdown experiments, we intranasally instilled mice with 50μg siRNA complexed with freshly extruded liposomes as previously described (Rowlands et al., 2011; Westphalen et al., 2014). Stock solutions of plasmids (2.5 μg/μl) were similarly complexed with freshly extruded unilamellar liposomes (20 μg/μl, 100 nm pore size) in sterile PBS to a final concentration of 1μg oligonucleotide/μl. Mice were intranasally instilled (50-75 μl) the nucleic acid-liposome mixture. Lungs from transfected animals were excised 48h later. Protein expression or knockdown was confirmed by immunoblotting.

#### Lentivirus cloning and production

From the *pMCUwt* plasmid, we PCR amplified the *cmv-MCU-GFP* sequence with following primers: 5’-TACGTAACCATGGCGGCCGCCGCAGGTAG-3’ and 5’-TCATTCCTTTTCTCCGATCTGTCGGAGGCTCGAG-3’. Thus, we introduced Snabl and Xhol restriction sites introduced in them. We used the PCR product for TA cloning. From the TA clone, the *cmv-MCU-GFP* sequence was excised using Snabl and Xhol. After the restriction digest, the excised sequence was ligated with the pGipz vector. The resulting clone was confirmed by restriction digest and sequencing. To make the virus, the lentiviral plasmid was co-transfected with envelop (pMD2.G) and packaging (pCMV-dR8.9) plasmids. Viral supernatants were harvested 48h after transfection, centrifuged, and filtered through a 0.45-mm filter (Millipore). Viral particles were pelleted at 50,000g x 2h at 4°C, resuspended in DMEM, aliquoted, and stored at −80°C. Viral titers were determined by Lenti-X qRT-PCR Titration Kit. Anesthetized animals were intranasally administered with 100μl viral suspension with a titer of 1×10^7^ pfu. Lungs were excised 21d after infection and MCU expression confirmed by immunoblotting and alveolar imaging.

#### Channelrhodopsin activation

To activate ChR2 in the alveolar epithelium, imaged alveolar fields were excited by mercury lamp illumination directed through an FITC (470±10 nm) filter. The duration of field excitation was 10-seconds.

#### Mitochondrial complex activity assay

We used a reported method to determine complex I-IV activities (Rera et al., 2011). Briefly, purified mitochondria were resuspended in complexes I-IV activity assay buffers. Activity of complexes I and II were measured as the rate of decrease in absorbance at 600nm. Whereas, activity of complexes III and IV were measured respectively, as the rate of increase and decrease in absorbance at 550nm. For each assay, absorbance was measured at the given wavelength every 10 sec for 30 min at 25°C using a SpectraMax microplate spectrophotometer. Enzymatic activities were normalized to protein concentrations. Complex I activity was calculated as the difference between activities measured when samples were incubated with 2μM Rotenone or ethanol.

#### Extravascular lung water

We determined blood-free lung water content by our reported method (Bhattacharya et al., 1989; Safdar et al., 2003). Briefly, we determined the wet and dry weights of excised lungs and quantified hemoglobin concentrations in lung homogenates to correct for blood water content in the wet/dry ratio.

#### Lung ATP determination

We used our reported colorimetric methods for lung ATP determination (Islam et al., 2012).

#### Bronchioalveolar lavage (BAL) neutrophil count

Exsanguinated animals were tracheally cannulated. BAL fluid was obtained by intratracheal instillations with 1 ml ice-cold sterile PBS. The recovered lavage fluid was centrifuged at 400g x 5 min at 4°C, and the pellet was resuspended in PBS supplemented with 1% BSA. Following erythrolysis, BAL cell numbers were counted using a Neubauer chamber. Neutrophils in the chamber were identified by fluorescence-tagged Ly-6G antibody.

### QUANTIFICATION AND STATISTICAL ANALYSIS

All major groups comprised a minimum of 3 mice each. Mean and SEM were calculated on a per-lung basis. We analyzed paired comparisons by the paired t-test and multiple comparisons by ANOVA with Bonferroni’s post-hoc analysis. Survival comparisons were analyzed by the Logrank test. All data are mean±SE. Significance was accepted at P<0.05.

---

**s-Figure 1.**
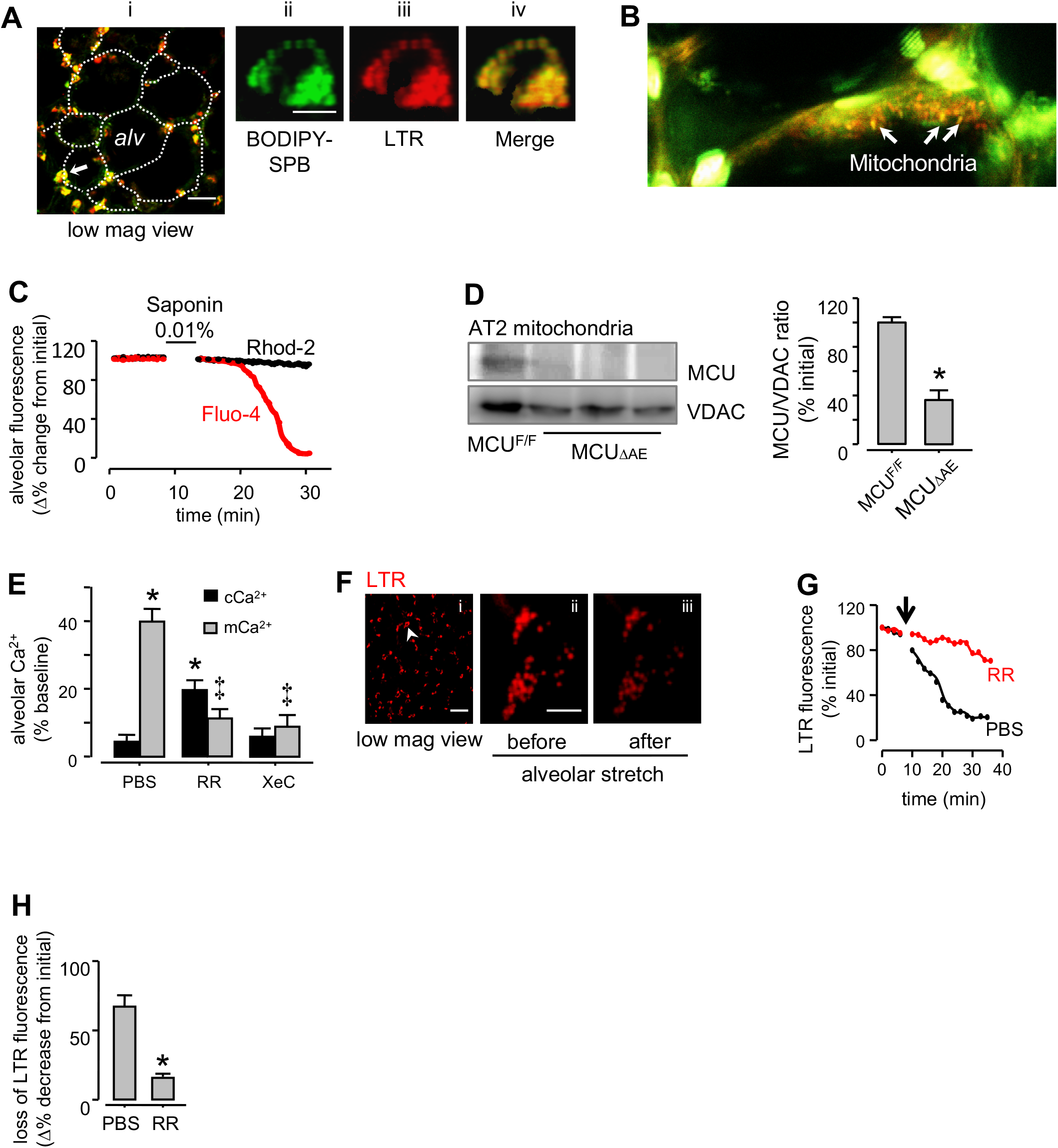
Mitochondrial assays in live alveoli. (A) Confocal images show AT2 *in situ* at low (i) and high (ii-iv) magnifications for the indicated region (*arrow in i*). Alveoli (*alv*) were stained with the lamellar body localizing dye, lysotracker red (*LTR*) and fluorescent surfactant protein B (*BODIPY-SPB*). Alveolar margins are delineated by dotted lines. Green and red channel renditions show individual lamellar bodies stained with SPB and LTR. Scale bars, 30μm (i) and 5μm (ii-iv). (B) Two-photon image (0.5 μm optical section) shows an alveolar septum with rhod-2 loaded mitochondria (arrows, *orange*), fluo-4 loaded cytosol (*green*) and cell nuclei (*white*). (C) Tracings show alveolar microinfusion of saponin selectively decreases the fluorescence of the cytosolic (fluo-4) but not the mitochondrial dye (rhod-2). (D) We crossed MCU floxed mice with mice expressing an inducible Cre driven by a surfactant protect C promoter (SPC-Cre). Pre-partum SPC-Cre induction by doxycycline caused epithelial MCU deletion in these mice (AE^MCU-^), as confirmed by MCU immunoblots in AT2 mitochondria derived from *MCU^F/F^* littermates and in alveolar MCU-null (AE^MCU-^) mice. Group data are quantification of MCU/VDAC band densities. *VDAC*, voltage dependent anion channel. n=3 mice for each bar, *p<0.05 versus MCU^F/F^. (E) Data are alveolar Ca^2+^ responses to alveolar stretch following indicated treatments. By microinfusion, alveoli were pre-treated with either *PBS*, Ruthenium Red (*RR*) or Xestospongin C (*XeC*) or. n= 4 lungs each group. *p<0.05 versus baseline and ‡p<0.05 versus corresponding PBS. (F-H) Images (*F*), tracings from a single experiment (*G*) and group data (*H*) show determinations of stretch induced surfactant secretion from intact alveoli of live lungs. Confocal image in low magnification (*i*) show an alveolar field stained with lamellar body localizing dye, lysotracker red (*LTR*). Magnification of a select cell (*arrow head in i*) show a single AT2 (*ii-iii*). Loss of LTR fluorescence following alveolar stretch (*arrow in G*) marks surfactant secretion. Indicated inhibitors were added by alveolar microinfusions. *RR*, ruthenium red. Scale bars, 10μm (i) and 5μm (ii-iii). n=4 lungs for each bar, *p<0.05 versus PBS. Group data are mean±SEM.

**s-Figure 2.**
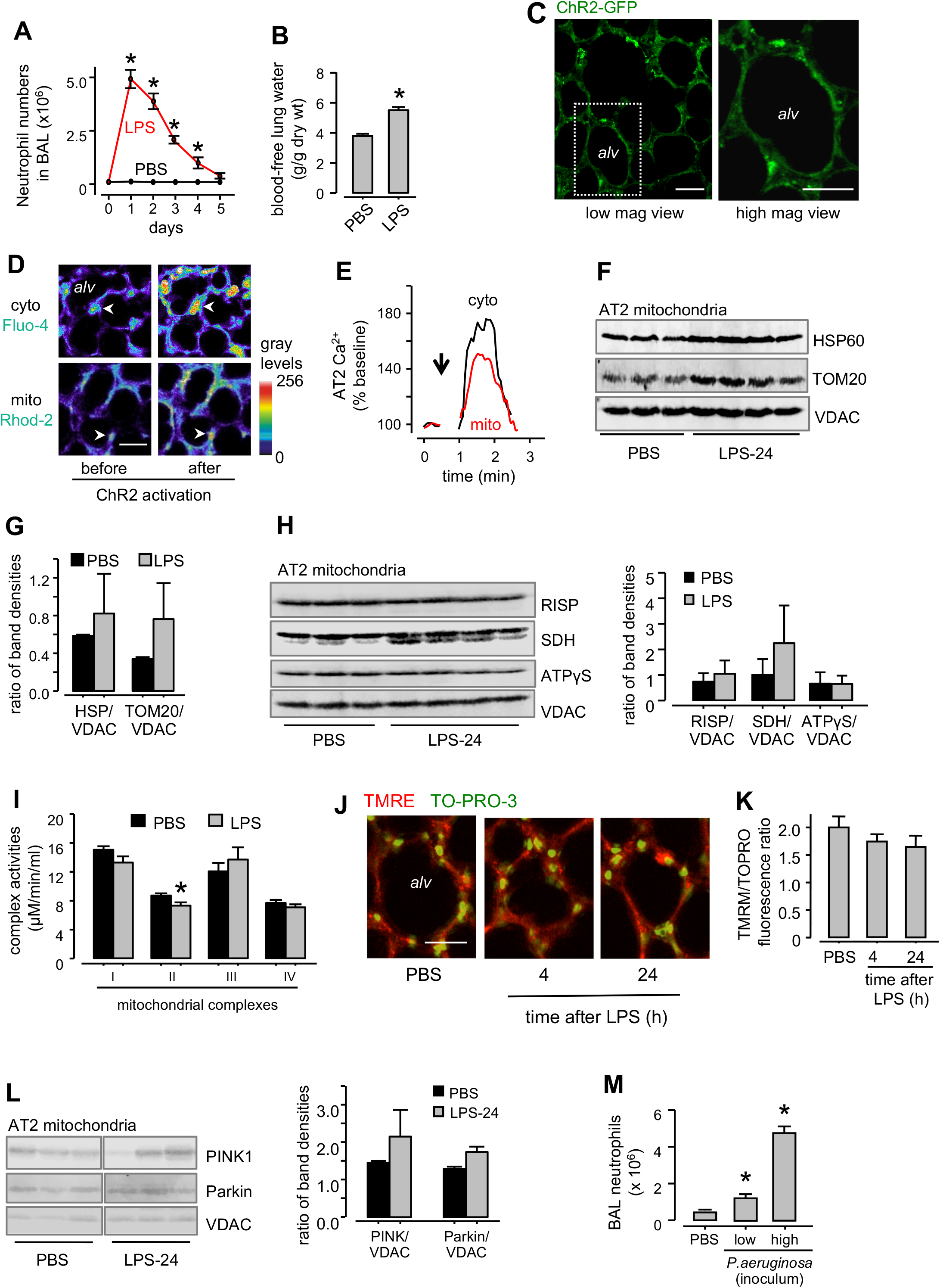
LPS depletes MCU. (A) Group data are determinations of neutrophil counts in bronchoalveolar lavage at indicated time points following intranasal instillations of LPS at a sublethal dose. n=4 mice per group, *p<0.05 versus corresponding PBS. (B) Determinations of blood-free lung water in mice 24h after indicated instillations. n=4 mice per group, *p<0.05 versus PBS. (C) Live confocal image in low magnification (*left*) shows GFP fluorescence of the expressed channelrhodopsin (*ChR2*) plasmid in an alveolar (*alv*) field. Magnification of the rectangle shows GFP expression in an alveolus (*right*). *FCK-ChR2-GFP* plasmid was intranasally instilled as a liposomal complex (75μg/mouse) and lungs excised 48h after plasmid instillation. Scale bars, 10μm. (D and E) Live confocal images in pseuodocolor rendition (*D*) and tracings from a single experiment (*E*) show simultaneous determination of AT2 (*arrowheads*) cytosolic (*cyto*) and mitochondrial (*mito*) Ca^2+^ responses to a single 10-second ChR2 activation (*arrow in E*). *Alv*, alveolus. Scale bar, 10μm. (F and G) Immunoblots for indicated proteins (*F*) and densitometry (*G*) are for AT2 mitochondria derived 24h after intranasal instillations of PBS or LPS at a sublethal dose. Protein densitometry is normalized for VDAC. *HSP60*, heat shock protein 60; *TOM20*, translocase of the outer membrane. n=4 lungs each bar. (H) Immunoblots and calculated band densities show expression of indicated protein in AT2 mitochondria 24h after intranasal instillations of PBS or sublethal LPS. *RISP*, Rieske’s ironsulfur protein; *SDH*, succinate dehydrogenase; *ATPγS*, adenosine trisphosphate gamma synthase. n= 4 lungs for group. *p<0.05 versus corresponding PBS. (I) Group data show determination of the activities of electron transfer complexes from alveolar mitochondria 24h after indicated instillations. LPS was instilled at a sublethal dose. n=4 lungs for each bar. *p<0.05 versus corresponding PBS. (J and K) Images (*J*) and group data (*K*) show alveolar determination of mitochondrial potential. Alveoli (*alv*) were stained with the potentiometric dye, tetramethylrhodamine ethyl ester *(TMRE)* and the nuclear-staining dye, TO-PRO-3. Scale bars, 10μm. n=4 lungs each bar. (L) In AT2 mitochondria, mitophagy proteins Pink1 and Parkin were immunoblotted and band densities calculated following indicated intranasal instillations. LPS was instilled at a sublethal dose. n= 4 lungs for each bar. *p<0.05 versus corresponding PBS. (M) Group data are determinations of neutrophil counts in bronchoalveolar lavage 24h after intranasal *P. aeruginosa* instillations. The low and high inoculum concentrations were 1×10^5^ and 1×10^6^ colony-forming units (CFU), respectively. n=4 mice per group, *p<0.05 versus corresponding PBS. Group data are mean± SEM.

**s-Figure 3.**
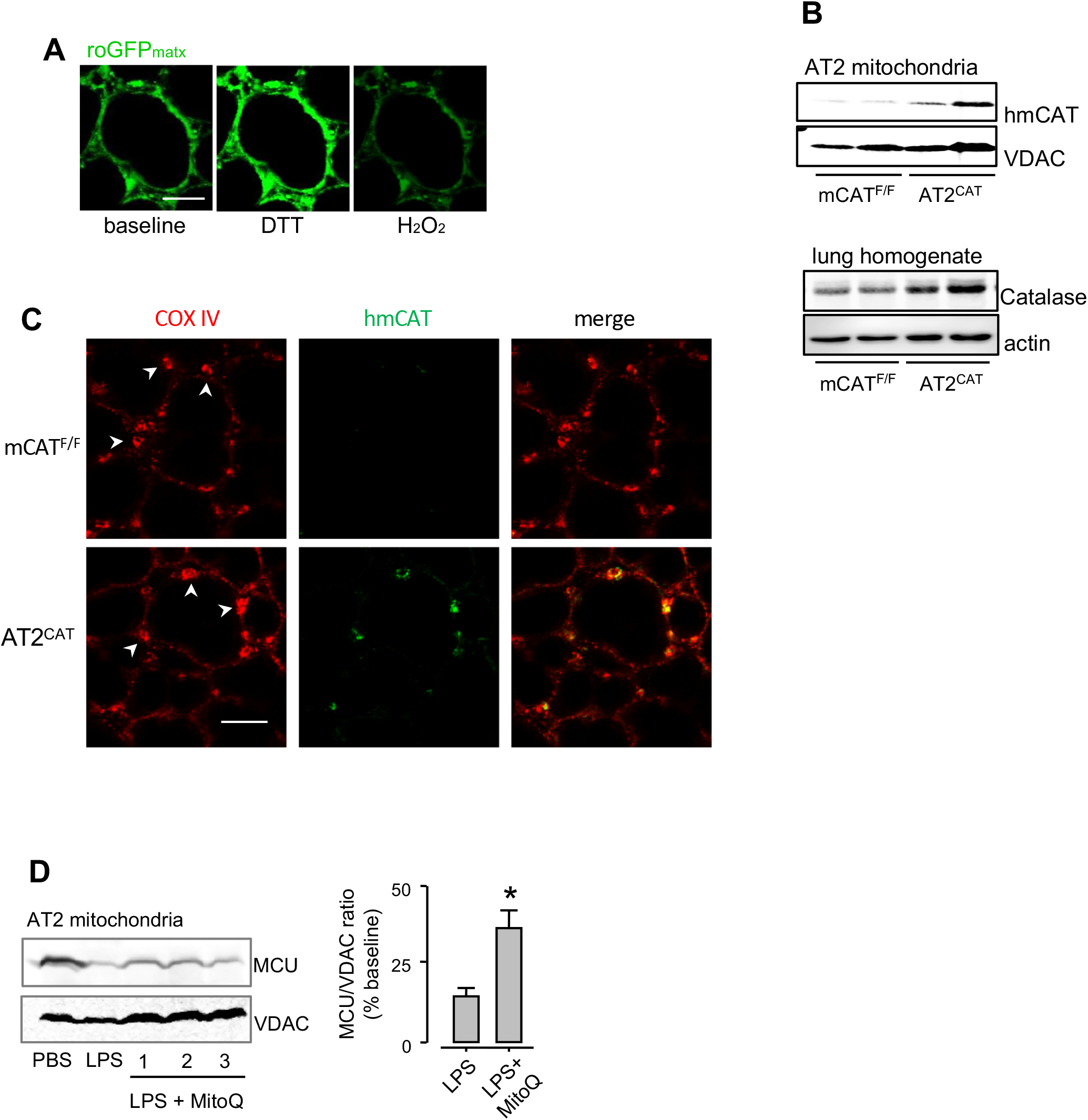
Inhibition of mitochondrial H_2_O_2_ blocks MCU depletion. (A) Confocal image sequence shows changes in mitochondrial matrix-targeted *roGFP* (*roGFPmatx*) fluorescence following indicated alveolar microinfusions. By alveolar microinfusion, the reducing agent, dithiothreitol (*DTT, 2mM*) and oxidizing agent, hydrogen peroxide (*H_2_O_2_, 100μM*) were added to obtain respectively, the maximum (fully reduced) and minimum (fully oxidized) fluorescence of the expressed probe. As a metric of mitochondrial H_2_O_2_, we determined roGFP oxidation (roGFPox) by the relation: roGFPox = (1-[F/F_DTT_]), where F and F_DTT_ are roGFP fluorescence values for the experiment and after exposure to the reducing agent, DTT. (B) Immunoblots for human catalase (*hmCAT*) in mitochondria derived from AT2 (*upper*) and total catalase in lung homogenates (*lower*). *mCAT^F/F^*, floxed mice for mitochondrial catalase (mCAT); *AT2^CAT^*, mice expressing catalase in alveolar mitochondria. (C) Images show *in situ* immunofluorescence of human catalase (*hmCAT, green*) and cytochrome oxidase IV (*COXIV, red*) in AT2 (arrowheads). Scale bar, 10μm. (D) MCU immunoblots in AT2 mitochondria (*left*) following indicated treatments. Mitochondria-targeted antioxidant MitoQ (mitoquinone mesylate, 100nM) was intranasally instilled 4h after intranasal instillation of LPS at a sublethal dose. Lungs were excised 24h after PBS or LPS instillations. Group data (*right*) are ratio of MCU and VDAC band intensities. n= 3 lungs each bar, *p<0.05 versus LPS alone. Group data are mean± SEM.

**s-Figure 4.**
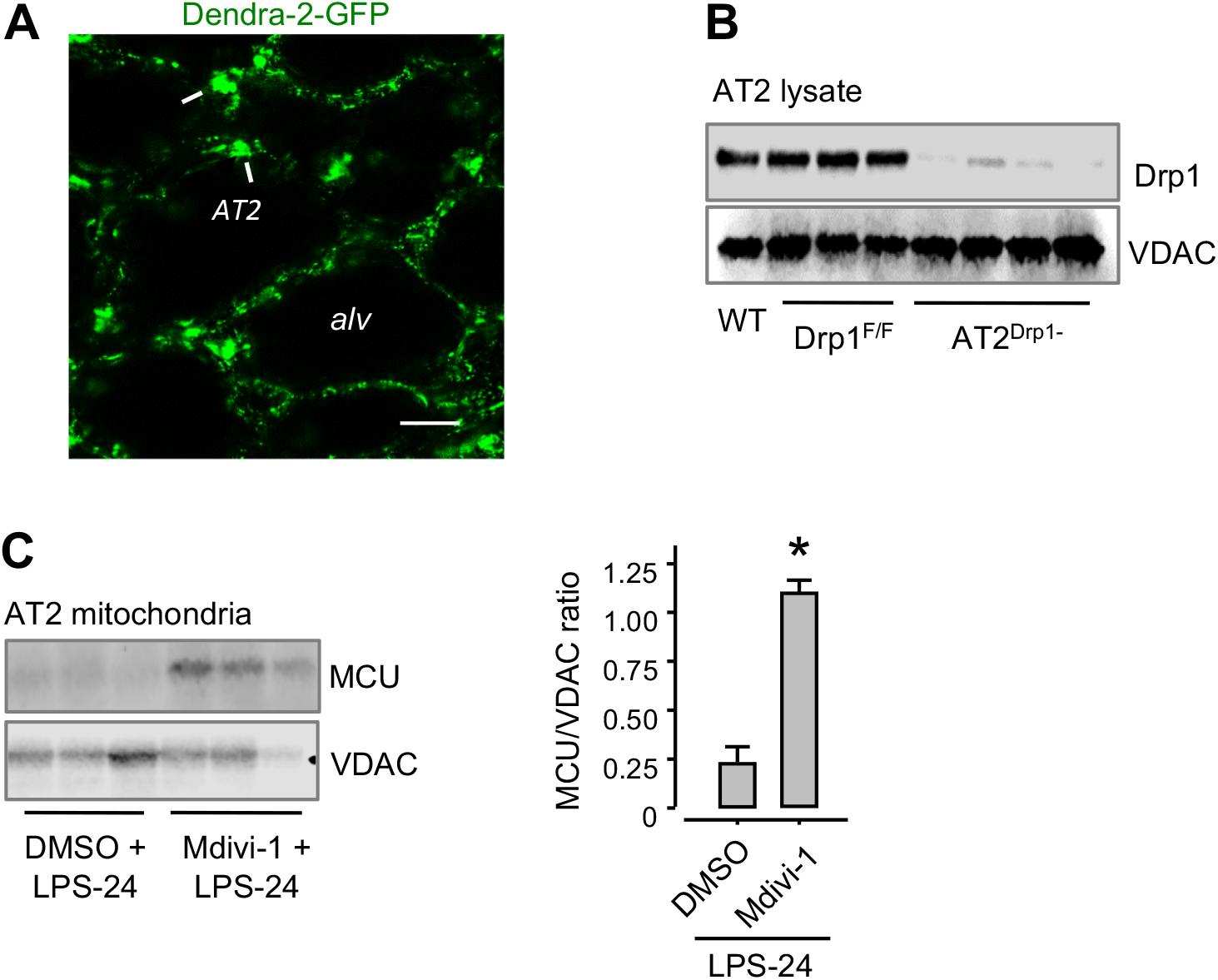
LPS causes mitochondrial redistribution. (A) Live confocal image shows alveolar (*alv*) fluorescence of the mitochondrial matrix-targeted protein, Dendra-2. Mitochondrial Dendra-2 expression was induced by breeding Pham floxed mice with E-2A cre mice. Expression of Dendra-2 is regulated by a cytochrome oxidase 8 (*COX8*) promoter. A high magnification image of a single AT2 is presented in Figure 4A. Scale bar, 5μm. (B) Drp1 immunoblots in AT2 homogenates derived from wild-type (WT), Drp1 floxed mice (*Drp1^F/F^*) or from Drp1 knockout mice in the AT2 (*AT2^DrP1-^*). We crossed Drp1^F/F^ mice with mice expressing an inducible Cre driven by a surfactant protect C promoter (SPC-Cre). Postpartum SPC-Cre induction by doxycycline caused epithelial Drp1 deletion in AT2 (AT2^Drp1-^). (C) MCU immunoblots and densitometry are for freshly isolated AT2 mitochondria following indicated treatments. Wild-type mice were intranasally instilled for 3 days at 12h intervals with 30 mg/kg of the selective Drp1 inhibitor, *Mdivi-1* or vehicle (*DMSO*, dimethyl sulfoxide) followed 24h later by intranasal instillations of LPS at a sublethal dose. Lungs were excised and AT2 mitochondria isolated 24h after LPS instillations. n=3 lungs for each bar. *p<0.05 versus corresponding DMSO. Group data are mean± SEM.

**s-Figure 5.**
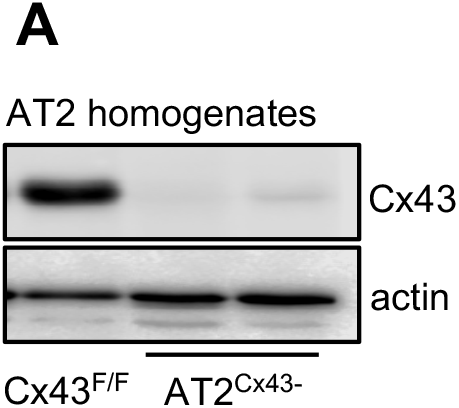
Generation of connexin 43 (Cx43) deleted mice in AT2. (A) We crossed connexin 43 (Cx43) floxed mice (Cx43^F/F^) with mice expressing an inducible Cre driven by a surfactant protect C promoter (SPC-Cre). Post-partum SPC-Cre induction by doxycycline caused AT2 Cx43 deletion in these mice (AT2^Cx43-^), as confirmed by Cx43 immunoblots in AT2 homogenates derived from mice of indicated genotype.

**s-Figure 6.**
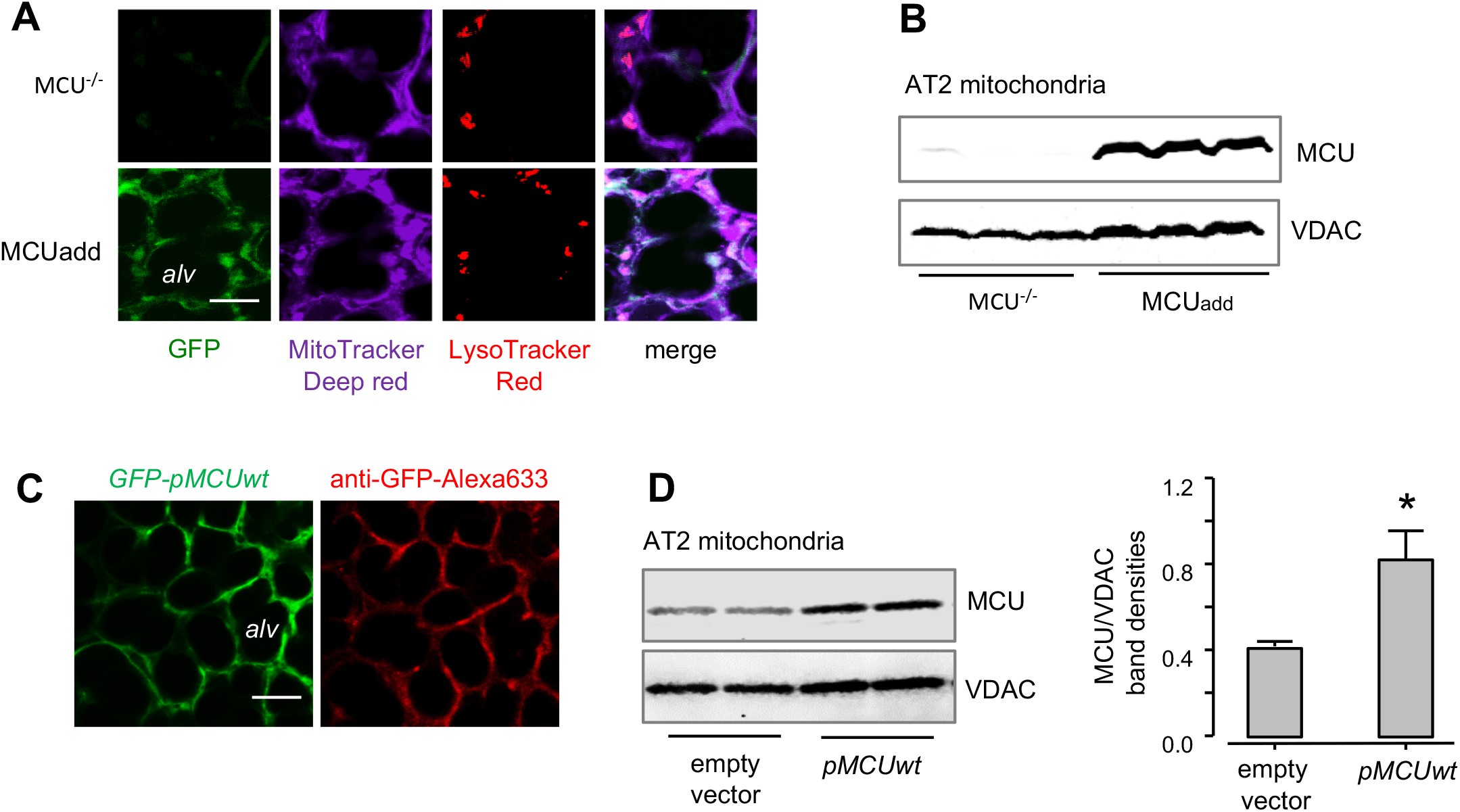
MCU overexpression in intact alveoli. (A and B) Live confocal images (*A*) and MCU immunoblots (*B*) show, respectively, alveolar *(alv)* fluorescence of the expressed MCU protein (GFP) and MCU protein in AT2 mitochondria derived from MCU-null (MCU^-/-^) or MCU^-/-^ with MCU add-back (MCUadd) mice. Epithelial mitochondria and AT2 were stained respectively, with alveolar microinfusions of mitotracker deep red (MTDR) and lysotracker red (LTR). Scale bar, 10μm. (C) Images show alveolar (alv) fluorescence of GFP-pMCUwt overexpression (left) and immunofluorescence of the expressed GFP (right). To elicit immunofluorescence, anti GFP mouse Ab tagged with AlexaFluor-633 (anti-GFP-Alexa633) was microinfused into fixed and permeabilized alveoli. *pMCUwt*, plasmid encoding for full length MCU. Scale bar, 10μm. (D) Immunoblots for MCU in AT2 mitochondria derived after intranasal instillations of empty vector and*pMCUwt*. Lungs were excised 48h after intranasal plasmid instillations. Group data are ratios of MCU/VDAC band densities. n=3 lungs each bar. Data are mean±SEM, *p<0.05 versus empty vector. Group data are mean ± SEM.

**s-Figure 7.**
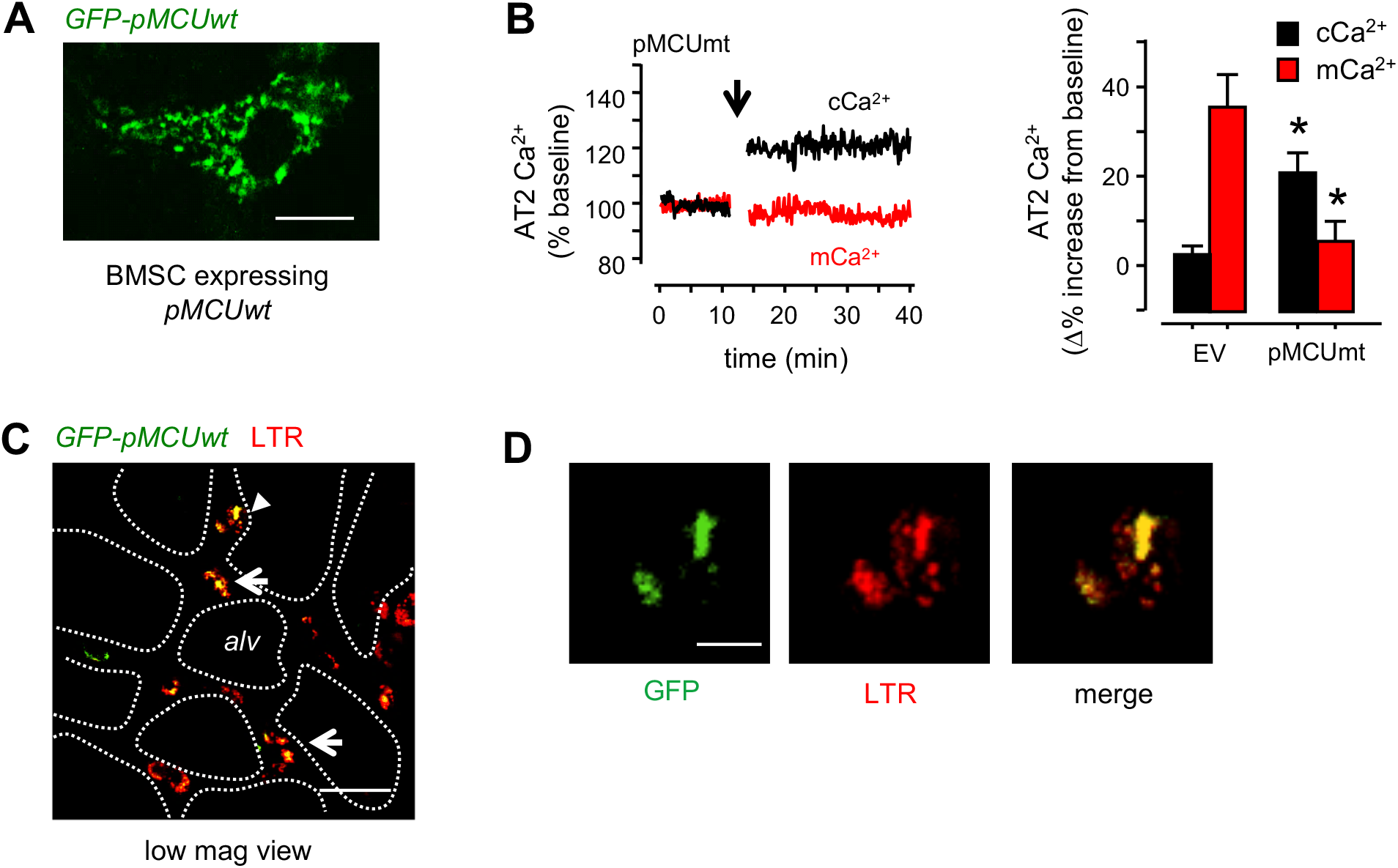
BMSCs expressing *pMCUwt* protect against LPS inflammation. (A) Live confocal image shows a single bone marrow-derived mesenchymal stromal cell (BMSC) expressing GFP-tagged *pMCUwt. pMCUwt*, plasmid encoding for full length MCU. Scale bar, 3 μm. (B) Tracings from a single experiment and group data show AT2 Ca^2+^ responses to alveolar stretch (arrow). Liposome-complexed pMCUmt plasmid (plasmid encoding for a mutant MCU) was given by intranasal instillation. Lungs were removed 48h after plasmid instillations. *EV*, empty vector. n=4 mice per group, *p<0.05 versus EV. (C) Live confocal image of an LPS-24 lung shows AT2 (LTR, *red*) and instilled BMSCs (arrows and arrowhead) expressing *GFP-pMCUwt* (*green*). Mice were given intranasal LPS instillations at a sublethal dose. BMSCs expressing *pMCUwt* were instilled 4h after LPS instillation. Lungs were excised and imaged 24h after LPS instillation. White lines represent alveolar (*alv*) margin. Scale bar, 15μm. (D) Magnified images of a single AT2 (*arrowhead in C*) show in green and red renditions respectively, fluorescence of BMSC-*pMCUwt* (*green*, left) and LTR (*red*, middle). Scale bar, 5μm. Group data are mean ± SEM.

**Supplementary Video 1. Alveolar stretch causes mCa^2+^ entry.** The movie is divided in three segments. In the first segment, image sequences show baseline fluorescence of a single AT2 (*cuboidal cell in the left*) in a live alveolus stained with the mitochondrial Ca^2+^-sensing dye, rhod-2. The second segment shows dark images corresponding to the alveolar stretch procedure. The third segment shows mages from the poststretch phase where rhod-2 fluorescence increases denoting increase of mitochondrial Ca^2+^.

## Notes

### Competing Interest Statement

The authors have declared no competing interest.

## REFERENCES

Ashino, Y., Ying, X., Dobbs, L.G., and Bhattacharya, J. (2000). [Ca(2+)](i) oscillations regulate type II cell exocytosis in the pulmonary alveolus. Am J Physiol Lung Cell Mol Physiol 279, L5–13.

Baughman, J.M., Perocchi, F., Girgis, H.S., Plovanich, M., Belcher-Timme, C.A., Sancak, Y., Bao, X.R., Strittmatter, L., Goldberger, O., Bogorad, R.L., et al. (2011). Integrative genomics identifies MCU as an essential component of the mitochondrial calcium uniporter. Nature 476, 341–345.

Bellani, G., Laffey, J.G., Pham, T., Fan, E., Brochard, L., Esteban, A., Gattinoni, L., van Haren, F., Larsson, A., McAuley, D.F., et al. (2016). Epidemiology, Patterns of Care, and Mortality for Patients With Acute Respiratory Distress Syndrome in Intensive Care Units in 50 Countries. JAMA 315, 788–800.

Bhattacharya, J., Cruz, T., Bhattacharya, S., and Bray, B.A. (1989). Hyaluronan affects extravascular water in lungs of unanesthetized rabbits. J Appl Physiol (1985) 66, 2595–2599.

Bhattacharya, J., and Matthay, M.A. (2013). Regulation and repair of the alveolar-capillary barrier in acute lung injury. Annu Rev Physiol 75, 593–615.

Boyden, E.S., Zhang, F., Bamberg, E., Nagel, G., and Deisseroth, K. (2005). Millisecond-timescale, genetically targeted optical control of neural activity. Nat Neurosci 8, 1263–1268.

Breslin, J.S., and Weaver, T.E. (1992). Binding, uptake, and localization of surfactant protein B in isolated rat alveolar type II cells. Am J Physiol 262, L699–707.

Carneiro, F.R.G., Lepelley, A., Seeley, J.J., Hayden, M.S., and Ghosh, S. (2018). An Essential Role for ECSIT in Mitochondrial Complex I Assembly and Mitophagy in Macrophages. Cell Rep 22, 2654–2666.

Cereghetti, G.M., Costa, V., and Scorrano, L. (2010). Inhibition of Drp1-dependent mitochondrial fragmentation and apoptosis by a polypeptide antagonist of calcineurin. Cell Death Differ 17, 1785–1794.

Cui, J., Li, Z., Zhuang, S., Qi, S., Li, L., Zhou, J., Zhang, W., and Zhao, Y. (2018). Melatonin alleviates inflammation-induced apoptosis in human umbilical vein endothelial cells via suppression of Ca(2+)-XO-ROS-Drp1-mitochondrial fission axis by activation of AMPK/SERCA2a pathway. Cell Stress Chaperones 23, 281–293.

Dai, D.F., Johnson, S.C., Villarin, J.J., Chin, M.T., Nieves-Cintron, M., Chen, T., Marcinek, D.J., Dorn, G.W., 2nd, Kang, Y.J., Prolla, T.A., et al. (2011). Mitochondrial oxidative stress mediates angiotensin II-induced cardiac hypertrophy and Galphaq overexpression-induced heart failure. Circ Res 108, 837–846.

De Stefani, D., Raffaello, A., Teardo, E., Szabo, I., and Rizzuto, R. (2011). A forty-kilodalton protein of the inner membrane is the mitochondrial calcium uniporter. Nature 476, 336–340.

Dominici, M., Le Blanc, K., Mueller, I., Slaper-Cortenbach, I., Marini, F., Krause, D., Deans, R., Keating, A., Prockop, D., and Horwitz, E. (2006). Minimal criteria for defining multipotent mesenchymal stromal cells. The International Society for Cellular Therapy position statement. Cytotherapy 8, 315–317.

Dong, Z., Shanmughapriya, S., Tomar, D., Siddiqui, N., Lynch, S., Nemani, N., Breves, S.L., Zhang, X., Tripathi, A., Palaniappan, P., et al. (2017). Mitochondrial Ca(2+) Uniporter Is a Mitochondrial Luminal Redox Sensor that Augments MCU Channel Activity. Mol Cell 65, 1014–1028 e1017.

Favaro, G., Romanello, V., Varanita, T., Andrea Desbats, M., Morbidoni, V., Tezze, C., Albiero, M., Canato, M., Gherardi, G., De Stefani, D., et al. (2019). DRP1-mediated mitochondrial shape controls calcium homeostasis and muscle mass. Nat Commun 10, 2576.

Georgiadou, E., Haythorne, E., Dickerson, M.T., Lopez-Noriega, L., Pullen, T.J., da Silva Xavier, G., Davis, S.P.X., Martinez-Sanchez, A., Semplici, F., Rizzuto, R., et al. (2020). The pore-forming subunit MCU of the mitochondrial Ca(2+) uniporter is required for normal glucose-stimulated insulin secretion in vitro and in vivo in mice. Diabetologia 63, 1368–1381.

Gherardi, G., Nogara, L., Ciciliot, S., Fadini, G.P., Blaauw, B., Braghetta, P., Bonaldo, P., De Stefani, D., Rizzuto, R., and Mammucari, C. (2019). Loss of mitochondrial calcium uniporter rewires skeletal muscle metabolism and substrate preference. Cell Death Differ 26, 362–381.

Gomes, L.C., and Scorrano, L. (2013). Mitochondrial morphology in mitophagy and macroautophagy. Biochim Biophys Acta 1833, 205–212.

Gu, L., Larson Casey, J.L., Andrabi, S.A., Lee, J.H., Meza-Perez, S., Randall, T.D., and Carter, A.B. (2019). Mitochondrial calcium uniporter regulates PGC-1alpha expression to mediate metabolic reprogramming in pulmonary fibrosis. Redox Biol 26, 101307.

Hough, R.F., Islam, M.N., Gusarova, G.A., Jin, G., Das, S., and Bhattacharya, J. (2019). Endothelial mitochondria determine rapid barrier failure in chemical lung injury. JCI Insight 4.

Ichimura, H., Parthasarathi, K., Lindert, J., and Bhattacharya, J. (2006). Lung surfactant secretion by interalveolar Ca2+ signaling. Am J Physiol Lung Cell Mol Physiol 291, L596–601.

Ichimura, H., Parthasarathi, K., Quadri, S., Issekutz, A.C., and Bhattacharya, J. (2003). Mechano-oxidative coupling by mitochondria induces proinflammatory responses in lung venular capillaries. J Clin Invest 111, 691–699.

Ip, W.K.E., Hoshi, N., Shouval, D.S., Snapper, S., and Medzhitov, R. (2017). Anti-inflammatory effect of IL-10 mediated by metabolic reprogramming of macrophages. Science 356, 513–519.

Ishihara, N., Nomura, M., Jofuku, A., Kato, H., Suzuki, S.O., Masuda, K., Otera, H., Nakanishi, Y., Nonaka, I., Goto, Y., et al. (2009). Mitochondrial fission factor Drp1 is essential for embryonic development and synapse formation in mice. Nat Cell Biol 11, 958–966.

Islam, M.N., Das, S.R., Emin, M.T., Wei, M., Sun, L., Westphalen, K., Rowlands, D.J., Quadri, S.K., Bhattacharya, S., and Bhattacharya, J. (2012). Mitochondrial transfer from bone-marrow-derived stromal cells to pulmonary alveoli protects against acute lung injury. Nat Med 18, 759–765.

Islam, M.N., Gusarova, G.A., Monma, E., Das, S.R., and Bhattacharya, J. (2014). F-actin scaffold stabilizes lamellar bodies during surfactant secretion. Am J Physiol Lung Cell Mol Physiol 306, L50–57.

Jain, P., Luo, Z.Q., and Blanke, S.R. (2011). Helicobacter pylori vacuolating cytotoxin A (VacA) engages the mitochondrial fission machinery to induce host cell death. Proc Natl Acad Sci U S A 108, 16032–16037.

Kamer, K.J., and Mootha, V.K. (2015). The molecular era of the mitochondrial calcium uniporter. Nat Rev Mol Cell Biol 16, 545–553.

Kelso, G.F., Porteous, C.M., Coulter, C.V., Hughes, G., Porteous, W.K., Ledgerwood, E.C., Smith, R.A., and Murphy, M.P. (2001). Selective targeting of a redox-active ubiquinone to mitochondria within cells: antioxidant and antiapoptotic properties. J Biol Chem 276, 4588–4596.

Kim, S.J., Syed, G.H., Khan, M., Chiu, W.W., Sohail, M.A., Gish, R.G., and Siddiqui, A. (2014). Hepatitis C virus triggers mitochondrial fission and attenuates apoptosis to promote viral persistence. Proc Natl Acad Sci U S A 111, 6413–6418.

Kopp, E., Medzhitov, R., Carothers, J., Xiao, C., Douglas, I., Janeway, C.A., and Ghosh, S. (1999). ECSIT is an evolutionarily conserved intermediate in the Toll/IL-1 signal transduction pathway. Genes Dev 13, 2059–2071.

Kwong, J.Q., Huo, J., Bround, M.J., Boyer, J.G., Schwanekamp, J.A., Ghazal, N., Maxwell, J.T., Jang, Y.C., Khuchua, Z., Shi, K., et al. (2018). The mitochondrial calcium uniporter underlies metabolic fuel preference in skeletal muscle. JCI Insight 3.

Kwong, J.Q., Lu, X., Correll, R.N., Schwanekamp, J.A., Vagnozzi, R.J., Sargent, M.A., York, A.J., Zhang, J., Bers, D.M., and Molkentin, J.D. (2015). The Mitochondrial Calcium Uniporter Selectively Matches Metabolic Output to Acute Contractile Stress in the Heart. Cell Rep 12, 15–22.

Lin, J.Y., Lin, M.Z., Steinbach, P., and Tsien, R.Y. (2009). Characterization of engineered channelrhodopsin variants with improved properties and kinetics. Biophys J 96, 1803–1814.

Lindert, J., Perlman, C.E., Parthasarathi, K., and Bhattacharya, J. (2007). Chloride-dependent secretion of alveolar wall liquid determined by optical-sectioning microscopy. American journal of respiratory cell and molecular biology 36, 688–696.

Lombardi, A.A., Gibb, A.A., Arif, E., Kolmetzky, D.W., Tomar, D., Luongo, T.S., Jadiya, P., Murray, E.K., Lorkiewicz, P.K., Hajnoczky, G., et al. (2019). Mitochondrial calcium exchange links metabolism with the epigenome to control cellular differentiation. Nat Commun 10, 4509.

Luchsinger, L.L., de Almeida, M.J., Corrigan, D.J., Mumau, M., and Snoeck, H.W. (2016). Mitofusin 2 maintains haematopoietic stem cells with extensive lymphoid potential. Nature 529, 528–531.

Luongo, T.S., Lambert, J.P., Yuan, A., Zhang, X., Gross, P., Song, J., Shanmughapriya, S., Gao, E., Jain, M., Houser, S.R., et al. (2015). The Mitochondrial Calcium Uniporter Matches Energetic Supply with Cardiac Workload during Stress and Modulates Permeability Transition. Cell Rep 12, 23–34.

Manczak, M., Kandimalla, R., Yin, X., and Reddy, P.H. (2019). Mitochondrial division inhibitor 1 reduces dynamin-related protein 1 and mitochondrial fission activity. Hum Mol Genet 28, 177–199.

Matsuda, N., Sato, S., Shiba, K., Okatsu, K., Saisho, K., Gautier, C.A., Sou, Y.S., Saiki, S., Kawajiri, S., Sato, F., et al. (2010). PINK1 stabilized by mitochondrial depolarization recruits Parkin to damaged mitochondria and activates latent Parkin for mitophagy. J Cell Biol 189, 211–221.

Mora, R., Arold, S., Marzan, Y., Suki, B., and Ingenito, E.P. (2000). Determinants of surfactant function in acute lung injury and early recovery. Am J Physiol Lung Cell Mol Physiol 279, L342–349.

Pan, X., Liu, J., Nguyen, T., Liu, C., Sun, J., Teng, Y., Fergusson, M.M., Rovira, II, Allen, M., Springer, D.A., et al. (2013). The physiological role of mitochondrial calcium revealed by mice lacking the mitochondrial calcium uniporter. Nat Cell Biol 15, 1464–1472.

Park, H.S., Liu, G., Liu, Q., and Zhou, Y. (2018). Swine Influenza Virus Induces RIPK1/DRP1-Mediated Interleukin-1 Beta Production. Viruses 10.

Perl, A.K., Wert, S.E., Nagy, A., Lobe, C.G., and Whitsett, J.A. (2002). Early restriction of peripheral and proximal cell lineages during formation of the lung. Proc Natl Acad Sci U S A 99, 10482–10487.

Polster, B.M., Nicholls, D.G., Ge, S.X., and Roelofs, B.A. (2014). Use of potentiometric fluorophores in the measurement of mitochondrial reactive oxygen species. Methods Enzymol 547, 225–250.

Preau, S., Delguste, F., Yu, Y., Remy-Jouet, I., Richard, V., Saulnier, F., Boulanger, E., and Neviere, R. (2016). Endotoxemia Engages the RhoA Kinase Pathway to Impair Cardiac Function By Altering Cytoskeleton, Mitochondrial Fission, and Autophagy. Antioxid Redox Signal 24, 529–542.

Raffaello, A., De Stefani, D., Sabbadin, D., Teardo, E., Merli, G., Picard, A., Checchetto, V., Moro, S., Szabo, I., and Rizzuto, R. (2013). The mitochondrial calcium uniporter is a multimer that can include a dominant-negative pore-forming subunit. EMBO J 32, 2362–2376.

Rasmussen, T.P., Wu, Y., Joiner, M.L., Koval, O.M., Wilson, N.R., Luczak, E.D., Wang, Q., Chen, B., Gao, Z., Zhu, Z., et al. (2015). Inhibition of MCU forces extramitochondrial adaptations governing physiological and pathological stress responses in heart. Proc Natl Acad Sci U S A 112, 9129–9134.

Rera, M., Bahadorani, S., Cho, J., Koehler, C.L., Ulgherait, M., Hur, J.H., Ansari, W.S., Lo, T., Jr., Jones, D.L., and Walker, D.W. (2011). Modulation of longevity and tissue homeostasis by the Drosophila PGC-1 homolog. Cell Metab 14, 623–634.

Rowlands, D.J., Islam, M.N., Das, S.R., Huertas, A., Quadri, S.K., Horiuchi, K., Inamdar, N., Emin, M.T., Lindert, J., Ten, V.S., et al. (2011). Activation of TNFR1 ectodomain shedding by mitochondrial Ca2+ determines the severity of inflammation in mouse lung microvessels. J Clin Invest 121, 1986–1999.

Safdar, Z., Wang, P., Ichimura, H., Issekutz, A.C., Quadri, S., and Bhattacharya, J. (2003). Hyperosmolarity enhances the lung capillary barrier. J Clin Invest 112, 1541–1549.

Schindelin, J., Arganda-Carreras, I., Frise, E., Kaynig, V., Longair, M., Pietzsch, T., Preibisch, S., Rueden, C., Saalfeld, S., Schmid, B., et al. (2012). Fiji: an open-source platform for biological-image analysis. Nat Methods 9, 676–682.

Schriner, S.E., Linford, N.J., Martin, G.M., Treuting, P., Ogburn, C.E., Emond, M., Coskun, P.E., Ladiges, W., Wolf, N., Van Remmen, H., et al. (2005). Extension of murine life span by overexpression of catalase targeted to mitochondria. Science 308, 1909–1911.

Shanmughapriya, S., Rajan, S., Hoffman, N.E., Zhang, X., Guo, S., Kolesar, J.E., Hines, K.J., Ragheb, J., Jog, N.R., Caricchio, R., et al. (2015). Ca2+ signals regulate mitochondrial metabolism by stimulating CREB-mediated expression of the mitochondrial Ca2+ uniporter gene MCU. Sci Signal 8, ra23.

Simula, L., Pacella, I., Colamatteo, A., Procaccini, C., Cancila, V., Bordi, M., Tregnago, C., Corrado, M., Pigazzi, M., Barnaba, V., et al. (2018). Drp1 Controls Effective T Cell Immune-Surveillance by Regulating T Cell Migration, Proliferation, and cMyc-Dependent Metabolic Reprogramming. Cell Rep 25, 3059–3073 e3010.

Smirnova, E., Griparic, L., Shurland, D.L., and van der Bliek, A.M. (2001). Dynamin-related protein Drp1 is required for mitochondrial division in mammalian cells. Mol Biol Cell 12, 2245–2256.

Smirnova, E., Shurland, D.L., Ryazantsev, S.N., and van der Bliek, A.M. (1998). A human dynamin-related protein controls the distribution of mitochondria. J Cell Biol 143, 351–358.

Steinhardt, R.A., Bi, G., and Alderton, J.M. (1994). Cell membrane resealing by a vesicular mechanism similar to neurotransmitter release. Science 263, 390–393.

Tomar, D., Jana, F., Dong, Z., Quinn, W.J., 3rd, Jadiya, P., Breves, S.L., Daw, C.C., Srikantan, S., Shanmughapriya, S., Nemani, N., et al. (2019). Blockade of MCU-Mediated Ca(2+) Uptake Perturbs Lipid Metabolism via PP4-Dependent AMPK Dephosphorylation. Cell Rep 26, 3709–3725 e3707.

Wang, P.M., Ashino, Y., Ichimura, H., and Bhattacharya, J. (2001). Rapid alveolar liquid removal by a novel convective mechanism. American journal of physiology. Lung cellular and molecular physiology 281, L1327–1334.

Wang, Y., Subramanian, M., Yurdagul, A., Jr., Barbosa-Lorenzi, V.C., Cai, B., de Juan-Sanz, J., Ryan, T.A., Nomura, M., Maxfield, F.R., and Tabas, I. (2017). Mitochondrial Fission Promotes the Continued Clearance of Apoptotic Cells by Macrophages. Cell 171, 331–345 e322.

Waypa, G.B., Marks, J.D., Guzy, R., Mungai, P.T., Schriewer, J., Dokic, D., and Schumacker, P.T. (2010). Hypoxia triggers subcellular compartmental redox signaling in vascular smooth muscle cells. Circ Res 106, 526–535.

Wert, S.E., Glasser, S.W., Korfhagen, T.R., and Whitsett, J.A. (1993). Transcriptional elements from the human SP-C gene direct expression in the primordial respiratory epithelium of transgenic mice. Dev Biol 156, 426–443.

West, A.P., Brodsky, I.E., Rahner, C., Woo, D.K., Erdjument-Bromage, H., Tempst, P., Walsh, M.C., Choi, Y., Shadel, G.S., and Ghosh, S. (2011). TLR signalling augments macrophage bactericidal activity through mitochondrial ROS. Nature 472, 476–480.

Westphalen, K., Gusarova, G.A., Islam, M.N., Subramanian, M., Cohen, T.S., Prince, A.S., and Bhattacharya, J. (2014). Sessile alveolar macrophages communicate with alveolar epithelium to modulate immunity. Nature 506, 503–506.

Wu, Y., Nguyen, T.L., and Perlman, C.E. (2020). Intravenous sulforhodamine B reduces alveolar surface tension, improves oxygenation and reduces ventilation injury in a respiratory distress model. J Appl Physiol (1985).

